# Small size of recorded neuronal structures confines the accuracy in direct axonal voltage measurements

**DOI:** 10.1101/2021.01.29.428709

**Authors:** Viktor János Oláh, Gergely Tarcsay, János Brunner

## Abstract

Patch-clamp instruments including amplifier circuits and pipettes affect the recorded voltage signals. We hypothesized that realistic and complete *in silico* representation of recording instruments together with detailed morphology and biophysics of small recorded structures will precisely reveal signal distortions and provides a tool that predicts native signals from distorted voltage recordings. Therefore, we built a model that was verified by small axonal recordings. The model accurately recreated actual action potential measurements with typical recording artefacts and predicted the native electrical behavior. The simulations verified that recording instruments substantially filter voltage recordings. Moreover, we revealed that instrumentation directly interferes with local signal generation depending on the size of the recorded structures, which complicates the interpretation of recordings from smaller structures, such as axons. However, our model offers a straightforward approach that predicts the native waveforms of fast voltage signals and the underlying conductances even from the smallest neuronal structures.

## Introduction

Patch-clamp technique is affected by limitations that originate primarily from the physical properties of the recording pipettes (Benndorf, 1995; Hamill, Marty, Neher, Sakmann, & Sigworth, 1981; Marty & Neher, 1995). Patch pipettes have significant resistance (R_pip_) and their glass wall represent a substantial capacitive surface (C_pip_) as well. Limitations can be reduced by optimizing the recording conditions (that is, with the reduction of the R_pip_ and C_pip_) and corrected by using compensatory mechanisms of the amplifiers. Under standard recording conditions in measurements from relatively large structures, such as neuronal somata, these optimizations and corrections can sufficiently reduce distortions to an acceptable level. Therefore, the difference between the recorded and native voltage signals are negligible during good current clamp conditions in most neuronal structures. However, the reduction of instrumental distortions could became inherently insufficient in cases where the recorded structures are small, such as most of the central synapses. Recording pipettes for small neuronal structures must have small tip, which inevitably results in larger R_pip_ values that substantial filters the recorded signals (Benndorf, 1995; Novak et al., 2013; Ying, Bruckbauer, Rothery, Korchev, & Klenerman, 2002). Consequently, the fast voltage signals such as the action potentials (APs) are particularly vulnerable to signal distortion associated with direct recordings in small axonal structures. The shape of axonal APs is a key determinant of neuronal signaling that affects neurotransmitter release and short-term dynamics in synaptic connections (Bean, 2007; Borst & Sakmann, 1999; Chao & Yang, 2019; Geiger & Jonas, 2000; Katz & Miledi, 1967; Kawaguchi & Sakaba, 2015; Zbili & Debanne, 2019). Therefore, accurate AP measurements are essential to understand fundamental mechanisms of the neuronal information flow. Recent developments of the recording apparatus allows collecting voltage signals from the finest axonal structures (Kawaguchi & Sakaba, 2015; Novak et al., 2013; Ritzau-Jost et al., 2021; Rowan, DelCanto, Jianqing, Kamasawa, & Christie, 2016; Vivekananda et al., 2017). However, the interpretation of these signals are still limited because signal distortions caused by the recording pipette and amplifier circuits remains elusive (Ritzau-Jost et al., 2021). We reasoned that as computational modeling allows the precise reconstruction of the native electrical behavior of the most complex neuronal structures (Henrik Alle, Roth, & Geiger, 2009; Beaulieu-Laroche & Harnett, 2018; Branco & Häusser, 2011; Jayant et al., 2017; Kwon, Sakamoto, Peterka, & Yuste, 2017) it should be similarly possible to simulate the behavior of the recording instruments.

With these in mind, we built and tested a realistic model that considers not only the biological structures but also amplifier and pipette features. With this complex model we simulated actual recording conditions with the aim of subsequent, simulated removal of the instrumental contributions and distortions from the recorded signals. Thus, this complex model allowed the correction of the distortions caused by patch-clamp recording instruments and predict isolated biological signals. We tested the model by predicting the native action potential waveform of a directly recorded small axonal varicosity. Our simulations showed that recording instrumentation not only filters the signal, but it directly interferes with native signal generation in small neuronal structures.

## Results

### In silico implementation of amplifier features

We developed and validated a realistic amplifier model working both in voltage clamp (VC) and current clamp (CC) mode using the NEURON simulation environment (Hines & Carnevale, 1997). The model needed to be suitable for both VC and CC modes, because instrumental compensations are typically determined in VC mode during seal formations and these settings are implemented for CC during voltage signals recordings. For precise implementation of the components of the amplifier circuit, we examined their capacitive and resistive properties using three test configurations (**Supplementary Figure 1**). The first test configuration was the isolated headstage in open circuit (test #1), which allowed the characterization of the high frequency boost unit (see below) and the capacitance compensation of the VC circuitry. Since open circuit measurements are not possible in CC mode, the circuit was closed via known resistors in the second test configuration (test #2). This configuration allowed the estimation of stray capacitance associated with the feedback resistor. The third test circuit was a modified model cell (test #3, type 1U, Molecular Devices), which represents whole-cell recording conditions as it includes an idealized cell and recording pipette with their resistive and capacitive components. In order to explore an extended range of signal processing capacity of the amplifier, model cell components were varied using custom capacitors and resistors. This test configuration allowed us to investigate the features of the capacitance neutralization of the CC circuitry.

The construction of our model amplifier initially based on idealized circuit representations, namely a resistive feedback circuit (a current-to-voltage converter) typical for the VC and a voltage follower circuit (a unity gain voltage buffer) with an idealized current source for the CC (Sherman-Gold, 2012; Sigworth, 1995; Wilson & Park, 1989) (**Figure 1**).

**Figure 1.**
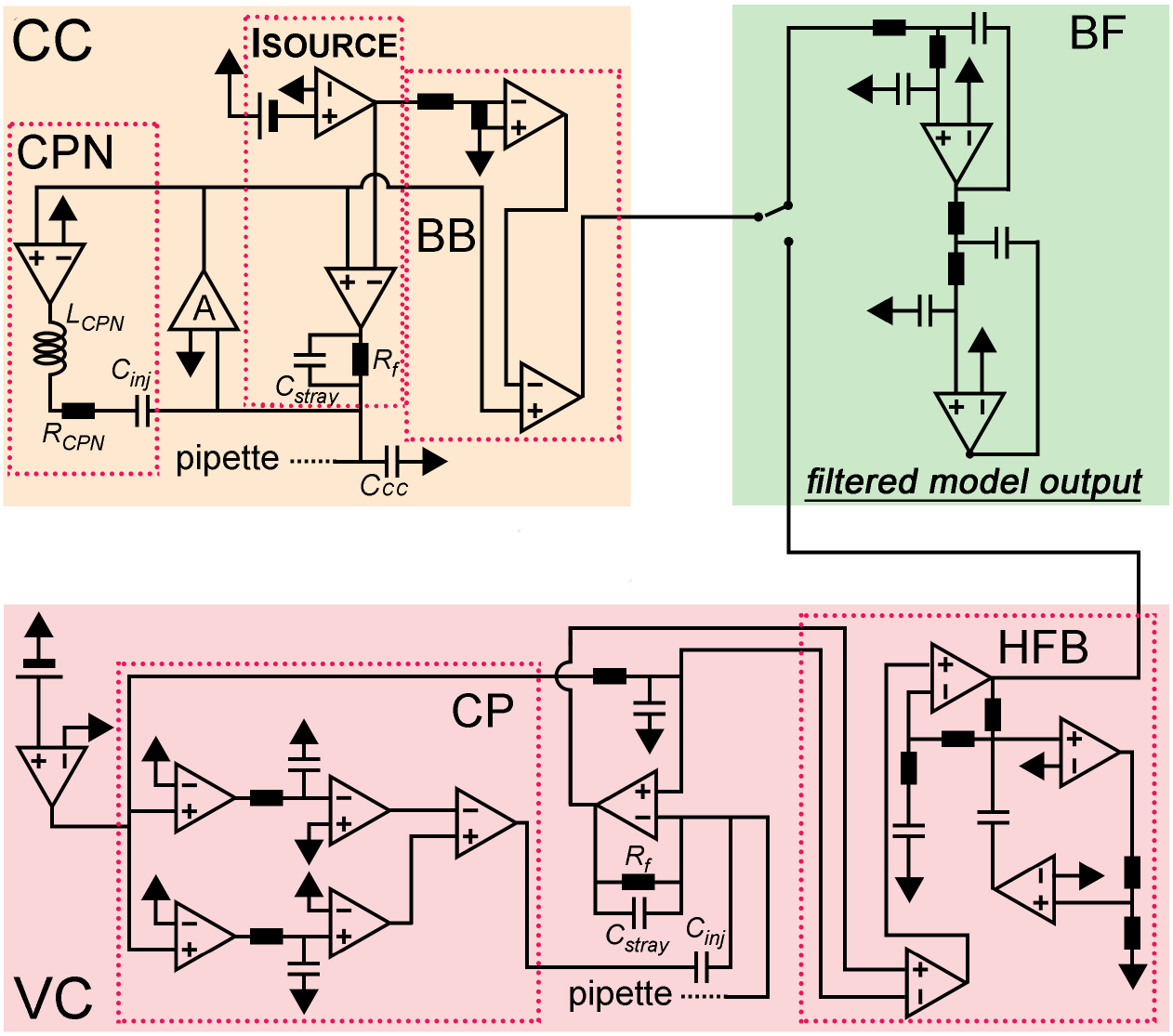
Model amplifier. Schematic circuit diagram of the model amplifier. The current clamp circuit (CC) consisted of a current source (Isource), an idealized voltage buffer (A), capacitance neutralization (CPN) and bridge balance (BB) circuits. The voltage clamp circuit (VC) consisted of a resistive feedback amplifier with dual capacitance compensation (CP) and the high frequency boost circuit (HFB). Outputs of the two amplifier modules are connected to a low-pass filter circuit (BF).

Operation of combined voltage- and current-clamp amplifiers requires a capacitor (C_inj_) for current injection in both VC and CC modes and a resistor (Rf) used as current generator in CC and as feedback resistor in VC (Strickholm, 1995) (**Figure 1**). To create realistic amplifier model, we first measured amplifier responses using the three test circuits, which were subsequently simulated allowing individual tuning of elementary circuit parameters. First, we measured C_inj_, assuming that the total capacitive load present in the isolated VC circuit (test #1) corresponds to the injector capacitor. By recording voltage-step-evoked capacitive current responses in open circuit configuration, we determined that the total capacitive load is 1.615 pF in our amplifier (MultiClamp700B). As virtually all real resistors, R_f_ has a certain amount of parasitic capacitance (C_stray_) which affects the performance characteristics of the amplifier. We next determined the size of this parasitic capacitance. In CC mode, the C_stray_ connected in parallel with the R_f_ (see Isource circuit on **Figure 1**) acts as a capacitance-neutralizing element. However, this neutralizing effect appears only when the input load is considerably smaller than the R_f_ which was set to 500 MΩ in the real amplifier (**Figure 2A left traces**). To characterize the size of C_stray_ associated with the R_f_, we adjusted the C_stray_ in the model to reproduce the evoked voltage responses recorded with the amplifier in test #2 configuration with 20, 50 and 100 MΩ input loads (**Figure 2A right traces**). These simulations revealed 0.38 pF C_stray_, thus we used this value in the model.

**Figure 2.**
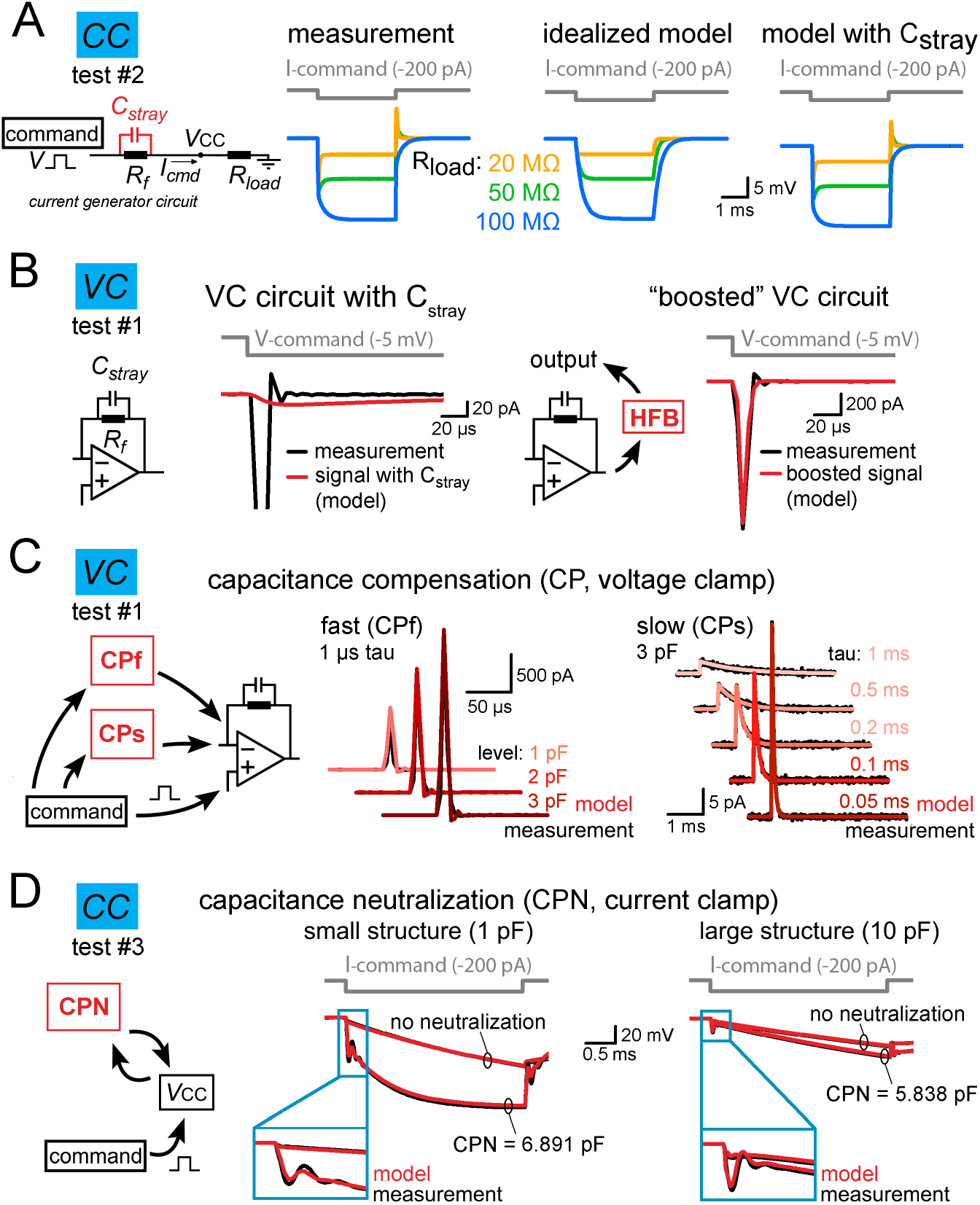
Implementation of individual amplifier components with realistic parameters. (A) The stray capacitance (C_stray_) associated with the current passing resistor (Rf) was predicted by simulating the voltage responses recorded in test #2 configuration. C_stray_ acts as a capacitance neutralizing element in CC mode. Because the capacitance neutralizing effect of C_stray_ depends on the load resistance (R_load_) attached to the amplifier input (V_CC_), we tested three scenarios using 20, 50 and 100 MΩ resistors. The left traces show measured voltage responses evoked by brief current injection in the presence of different R_load_. Idealized (i.e. C_stray_ free) model responses are shown in the middle for comparison. Right, model responses with 0.38 pF C_stray_ replicated the observed amplifier behavior. (B) C_stray_ of R_f_ slows down the current responses in the VC model (red trace, left panel). This effect is compensated in the actual amplifiers by using high frequency boost circuit (HFB). The implementation of a high frequency boost circuit in the model restored the response speed (red trace, right panel). (C) Capacitance compensation in the VC model (red) faithfully replicates the magnitude and the time course of amplifier-generated compensatory currents (black, test circuit#1) for both the fast (CPf) and slow (CPs) compensations. (D) Comparison of simulated (red) and recorded (black) voltage signals obtained with different capacitance neutralization (CPN) in CC mode from a small structure (middle panel) and from a large structure (left panel). Notice that the instrumental CC model faithfully replicates the neutralizing capability of the amplifier and the neutralization associated signal artefacts regardless of the applied CPN settings.

Next, we added the remaining amplifier components to the model one-by-one. First, we focused on the VC operations whose speed depends on a dedicated high frequency boost circuit (Sigworth, 1995). Amplifier are built with this compensation mechanism because C_stray_ substantially reduces the output bandwidth of the feedback circuit in VC mode (time constant of the capacitive relaxation with the previously determined 0.38 pF C_stray_: 191.92 μs, **Figure 2B left**). We added a simplified boosting unit tuned to accelerate model responses to the experimentally observed amplifier speed (3.85 μs vs. 3.19 μs, real vs. boosted model, **Figures 1 and 2B right**). Next, we implemented two pipette capacitance compensation circuits in the VC model. Fast capacitance compensation (CPf, 0-16 pF, 0.5-1.8 μs) cancels the majority of C_pip_-induced current transients, while the slow capacitance compensation (CPs, 0-3 pF, 10-4000 μs) reduces the slower instrumental capacitive components (Sigworth, 1995) (not equivalent to whole-cell compensation that was not implemented in this model **Figures 1 and 2C**).

We also implemented two compensatory mechanisms for the CC mode, namely the capacitance neutralization (CPN) and the bridge balance (BB) compensation (**Figure 1**). The CPN circuit is a positive feedback loop that feeds the pipette voltage back to the input through the C_inj_ in order to discharge the pipette capacitance (Sherman-Gold, 2012; Wilson & Park, 1989). Due to this positive feedback, the recordings are prone to oscillate as CPN level approaches a fully compensated state. Such evolving oscillation carry information about the intrinsic behavior of the CPN circuit. Therefore, the oscillating signal (dampening and frequency profile) can be employed for determining a minimal set of passive circuit elements necessary the re-create the CPN behavior. Specifically, we measured the maximal possible capacitance neutralization where the recording is still stable using test#3 circuit. Pipette parameters (10MΩ, 2.8 pF) and the cell-equivalent resistor (500 MΩ) was fixed and only the capacitance of the cell was varied from 0.75 pF to 46.7 pF. The model reproduced the recorded signal artefacts when a resistor (1.49 MΩ) and an inductor (18.3 H) was incorporated to the CPN circuit in series with the C_inj_ (**Figures 1 and 2D**). Amplifiers are typically supplied with BB compensatory mechanism to eliminate the voltage drop across the access resistance (Araki & Otani, 1955; Sherman-Gold, 2012). In the model, we subtracted a scaled version of the command signal from the recorded voltage (**Figure 1**). We also extended our model with a four-pole low-pass Bessel filter unit with adjustable cutoff frequencies from 0.5 kHz to 100 kHz (**Figure 1**).

Altogether, by using a measurement-based approach we created an amplifier model, in which both the VC and CC operation and their specific compensatory capabilities show realistic behavior.

### Pipette implementation considers the observed nonuniform C_pip_ and R_pip_ distributions

Next, we focused on the accurate implementation of patch pipettes. To properly characterize the distribution of resistance along patch pipettes that are suitable for recordings from small axons, we repeatedly broke small pieces from the end of the pipettes and determined the resistance as a function of tip distance (**Figure 3A**, n = 55 resistance measurement). In agreement with the theoretical considerations (Benndorf, 1995; Ying et al., 2002), we found that pipette resistance drops sharply after the tip, falling below one MΩ within the first millimeter (0.33 ± 0.18 MΩ for measurements with tip distance over 1 mm, n=20 measurement). To determine the capacitance distribution of our pipettes we systematically varied the immersed length of the pipettes in the recording solution and measured the capacitance, which corresponds to the cumulative capacitance of the dipped part (Cornwall & Thomas, 1981) (**Figure 3A**, n = 656 capacitance measurement). The cumulative capacitance increased within the first 3 mm from the tip, whereas the remaining part of the pipette had only moderate contribution to the total capacitance suggesting inhomogeneous capacitance distribution along pipettes (0.03 ± 0.07 pF, 2.52 ± 0.3 pF and 3.1 ± 0.25 pF at 0.1 mm, 3 mm and 7 mm tip distances respectively, n=39, 40 and 38 measurements, **Figure 3A**). A key parameter that defines the pipette capacitance is the ratio of the outer and inner pipette diameters (R_OI_) (Benndorf, 1995; Cornwall & Thomas, 1981). One potential explanation for the inhomogeneous capacitance distribution is that the R_OI_ is not constant along the pipette but it decreases toward the tip. To test this possibility and verify the predictions of the capacitance distribution measurement, we directly measured the R_OI_ of the recording pipettes. In order to precisely measure the edge of the pipette walls and avoid optical distortions by the curved glass walls, we carefully split the pipettes using a custom-built grinding system and measured their inner and outer diameters along the longitudinal axis (**Figure 3B**, see Methods section for details). Consistent with the dipping measurements, R_OI_ decreased toward the tip (R_OI_ <2mm = 1.6 ± 0.02 vs. R_OI_ >5mm = 2.04 ± 0.02, n = 4 pipettes) explaining the larger contribution of the tip region to the total pipette capacitance.

**Figure 3.**
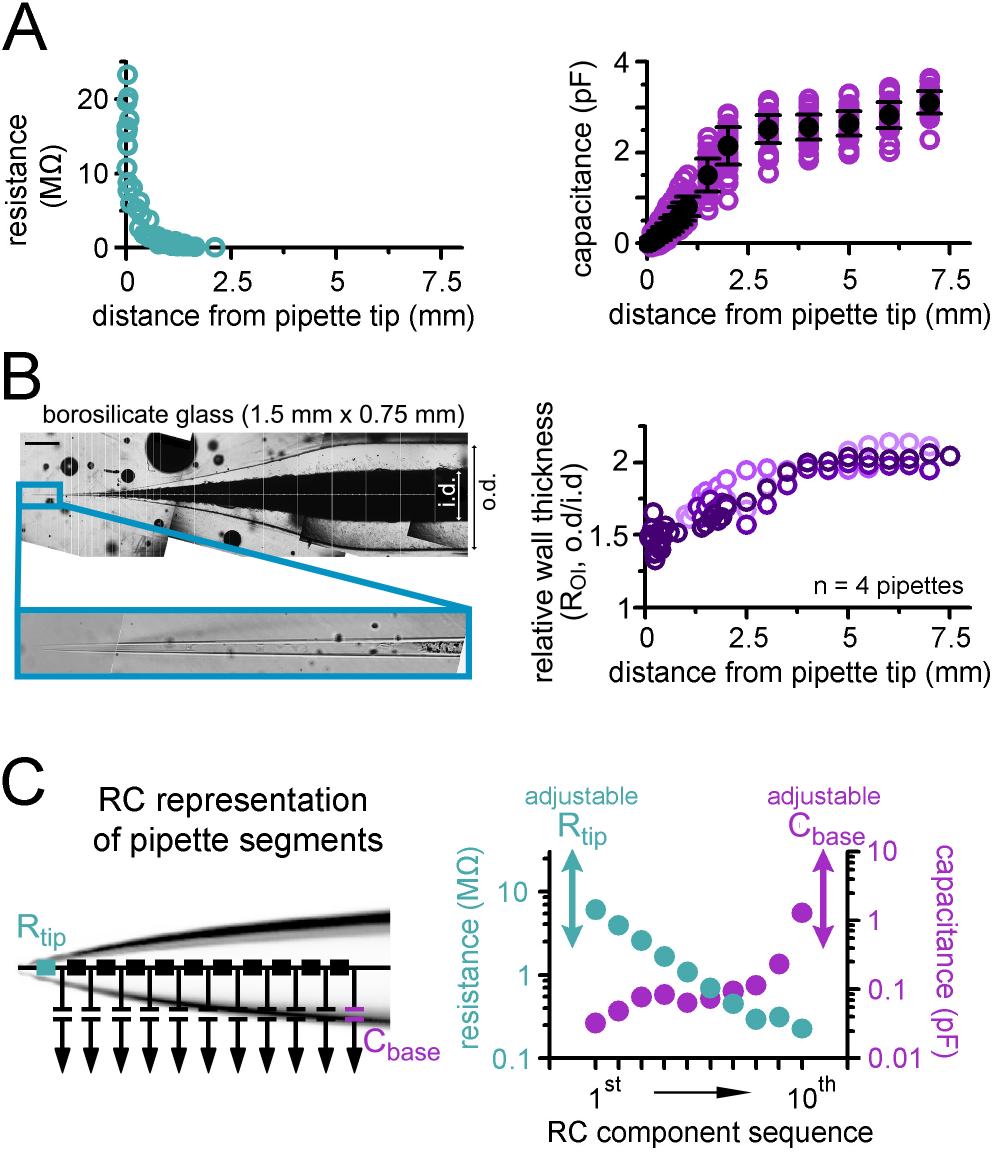
Implementation of pipettes with non-uniform C_pip_ and R_pip_ distributions. (A) Pooled data of R_pip_ (left graph) and C_pip_ (right graph) as a function of tip distance. (B) A representative imaging plane that was used for the measurement of the relative wall-thickness (R_oi_) that is, the ratio of outer and inner diameter (o.d. and i.d., respectively) of the pipettes. Scale bar: 0.5 mm. Graph on the right shows that the wall of the pipettes is much thinner toward the tips (i.e. the inner diameter is larger than predicted from the R_oi_ of the original glass; the 4 measured pipettes are shown in different shades of purple). (C) Pipettes were implemented as 10 independent RC units, a resistor (R_tip_, on the left) and a capacitor (C_base_, on the right). The two latter components allow the adjustment of the model to fit the differences of R_pip_ and C_pip_ of individual pipettes. Graph on the right indicates the actual capacitance (purple) and resistance (green) values of the 10 fixed RC motifs.

Based on these measurements, we created the skeleton of a “prototypical” patch pipette model from 10 RC units to consider the inhomogeneous distribution of capacitance and resistance (**Figure 3C**). Model pipette parameters can be adjusted to account for variability across individual pipettes using an additional resistor placed to the tip and a capacitor placed to the back to tune the R_pip_ and C_pip_ of individual pipettes.

### Reconstitution of the native electrical behaviour of a small axon

To test the efficacy of the complex instrumental model in predicting undisturbed fast neuronal membrane responses from distorted recordings, we used patch-clamp data from a small (<1 μm) *en passant* axonal varicosity of an identified hippocampal mossy fiber axon (MF, **Figure 4A**). Recordings from submicron-sized neuronal structures are supposedly substantially distorted as they can be patched only with high access resistance and their capacitance is in the range of the remaining uncompensated instrumental capacitance.

**Figure 4.**
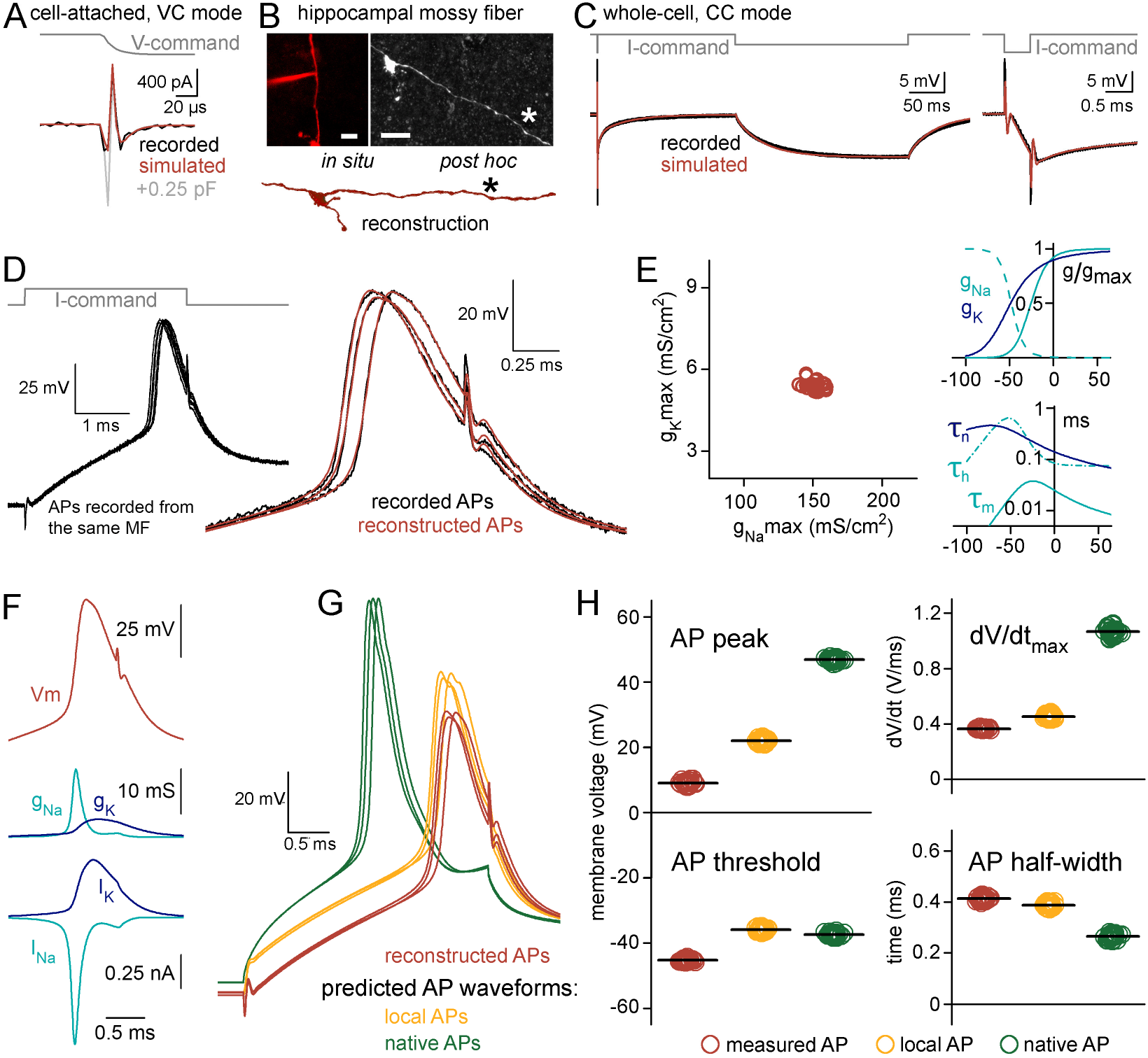
Reconstitution of the undisturbed membrane dynamics of a recorded axon. (A) Current responses to −20 mV voltage steps recorded from the axonal membrane in on-cell mode (black) and in the model with 7.097 pF (red) and 7.347 pF (grey) C_tot_. Notice the sensitivity of simulated responses to small differences in the instrumental capacitance applied in the model. (B) Confocal z-stack images show the recorded axon at the end of the experiment (left) and after the anatomical recovery (right). Bottom, part of the reconstructed morphology. Asterisks mark the recording position. Scale bars: 10 μm (C) Voltage responses in the recorded axon (black) and in the passive cable model with the added instrument (red) to short (3 ms, −50 pA) and long (250 ms, −2.5 pA) current stimuli. The short pulse is shown at an expanded timescale on the right. (D) Recorded (black) and simulated APs (red) evoked by brief (3 ms, 86 pA) current stimuli. APs were simulated in the complex model that included the instrument, the passive cable and Hodgkin-Huxley type sodium and potassium conductances. (E) Left, distribution of optimal maximal Na^+^ (gNa_max_) and K^+^ (gK_max_) conductance densities obtained from 30 recorded and independently fitted APs. Right, averaged gating properties and voltage dependence of the conductances that were resulted in by the fits that recreated the recorded APs. (F) Simulated AP waveform (top) with the underlying conductances (middle, g_Na_ and g_K_) and the modeled ionic currents (bottom, I_Na_ and I_K_). (G) The simulations with the confirmed conductance sets allowed to see the waveforms of the same APs not only within the pipette (red, corresponding to measured APs) but also the simultaneous AP waveforms within the axon (orange, corresponding to local APs) and, after the removal of the recording instrument from the model, the native, undisturbed AP waveforms (green). (H) Differences in waveform parameters of measured, local and native APs. Individual points show the peak, threshold, maximum rate of the rise (dV/dt_max_) and half-width of 30 independently simulated APs in the three points of view. Horizontal black lines indicate mean values.

The experimental conditions needed to be realistically modeled, including the precise morphology and electrical properties of the biological structure and the instrumental conditions (**Supplementary Figure 2**, see Methods for details). First, we characterized the total instrumental capacitance (C_tot_, the sum of all amplifier-, holder- and pipette-related capacitances) present in the actual recording (**Figure 4A**, C_tot_ : 7.097 pF from which 3.23 pF is the C_pip_ and 2 pF is the capacitance of the pipette holder). Target trace for this estimation was recorded using both fast and slow pipette capacitance compensation of the amplifier with the highest frequency resolution (output filter was bypassed) and we set the model accordingly.

Next, we precisely reconstructed the morphology of the biocytin-labelled and fluorescently recovered MF axon because signal propagation strongly depends on the length, diameter and their inhomogeneties of the biological structures (Goldstein & Rall, 1974; Manor, Koch, & Segev, 1991) (**Figure 4B**). As it is characteristic for hippocampal mossy fibers the 519 μm long reconstruction of the recorded MF included large (>3 μm) terminals, filopodial extensions and small (<1.5 μm) *en passant* varicosities within the *stratum lucidum* of the CA3 area (Acsady, Kamondi, Sik, Freund, & Buzsáki, 1998; Rollenhagen et al., 2007).

We next determined the cable properties; specific membrane resistance (R_m_), membrane capacitance (C_m_), and intracellular resistivity (R_i_) of the particular axon. For this, we optimized the model to the experimentally recorded voltage responses evoked by short and long current stimuli (**Figure 4C**) (Nörenberg, Hu, Vida, Bartos, & Jonas, 2010; Roth & Häusser, 2001; Schmidt-Hieber, Jonas, & Bischofberger, 2007; Szoboszlay et al., 2016). The advantages of the complex (instrumental + biological) model became obvious in these fittings. Pipette artefacts can markedly contaminate the onset of the evoked responses upon current injection (Major, Larkman, Jonas, Sakmann, & Jack, 1994). The incorporation of the complete experimental instrument to the model precisely reproduced these stimulus artefacts, therefore, allowing us to isolate biological contributions and obtain the passive cellular parameters. The predicted cellular parameters (R_m_ = 60.0 kΩ*cm^2^, a C_m_ = 0.65 μF*cm^−2^ and R_i_ = 147.3 Ω*cm) match with the data reported for MFs using recordings from large terminals (S. Hallermann, Pawlu, Jonas, & Heckmann, 2003). It is important to note that the similarities of the simulated and recorded fast voltage transients further verify our complex model.

Next, we simulated the active ionic mechanisms underlying the recorded AP waveforms. APs were evoked with brief current stimuli (**Figure 4D and E**, 3 ms, 86 pA). To simulate APs we tuned Hodgkin-Huxley-type sodium and potassium conductances (Hodgkin & Huxley, 1952) (modified version of the built-in mechanism in the NEURON simulation environment). Optimization of the density, kinetics and voltage dependence resulted in AP waveforms closely matching to the recorded ones (absolute AP peak: 9.0 ± 0.12 mV vs. 9.28 ± 0.12 mV, recorded vs. simulated APs, respectively, AP half-width: 0.52 ± 0.002 ms vs. 0.52 ± 0.002 ms,, AP threshold: −45.28 ± 0.07 mV vs. −44.27 ± 0.16 mV, maximum rate of rise: 363.26 ± 0.93 vs. 432.27 ± 3.56, n = 30 APs; **Figure 4D**). This suggest that the complex instrumental model in combination with traditional conductance functions is sufficient to reconstruct the recorded AP waveforms from potentially distorted recordings. Importantly, although each of the target APs was fitted independently, the optimal model conductance parameters were confined to a narrow range within the parameter space (coefficient of variation = 3% for both the Na^+^ and K^+^ conductance predictions, maximal g_Na_ density: 151.7 ± 5.23 mS/cm^2^, maximal g_K_ density: 5.41 ± 0.16 mS/cm^2^, n = 30; **Figure 4E**) indicating that the optimization provided a unique solution for the experimental data. Analysis of the error between the fit and its target data in different Na^+^ and K^+^ conductance combinations also revealed a single minimum that coincided with the best fit parameter combinations (**Supplementary Figure 3**). Furthermore, the optimal conductance parameters (i.e. their maximal conductance levels, their voltage dependence, and their activation and inactivation time constants), as well as the resulted ionic currents, are similar to previously described mechanisms underlying cortical axonal APs at near-physiological temperature (Henrik Alle et al., 2009; Geiger & Jonas, 2000; Stefan Hallermann, De Kock, Stuart, & Kole, 2012; Schmidt-Hieber & Bischofberger, 2010) (H. Alle, Kubota, & Geiger, 2011; Engel & Jonas, 2005; Hu & Jonas, 2014) (**Figure 4 E and F**), confirming the validity of our AP-reconstitution approach.

The relatively high series resistance (R_access_) in the recording (modeled R_access_: 53.2 ± 1.02 MΩ in the AP reconstitution simulations) can result in significant pipette filtering. The complex model, however, allowed us to investigate not only the APs recorded through the pipette but also the local spike that occurred within the axon while it was patched (**Figure 4G**). These local APs are not affected by the filtering effect caused by the pipette, so we could directly quantify filtering effects by comparing the *local* and *recorded* axonal AP waveforms (**Figure 4H**). Because of the filtering, local spikes had larger peak amplitudes and faster time course than the recorded APs (absolute peak: 21.95 ± 0.12 mV vs. 9.0 ± 0.12 mV, local vs. measured AP, respectively, *p = 6.88*10^−60^, t(29) = −552.75 paired sample t-test*, n = 30 APs, half-width: 0.48 ± 0.002 ms vs. 0.52 ± 0.002 ms, local vs. measured AP, respectively, *p = 1.99 * 10^−17^, t(29) = 18.23, paired sample t-test*, n = 30 APs, **Figure 4H**). As expected, the maximal rate of rise (dV/dt_max_) during the upstroke of spikes was the most different between local and measured APs as it is the most sensitive to low-pass filtering introduced by the pipette (453.11± 2.54 V/s vs. 363.26 ± 0.93 V/s, respectively, *p=4.69 * 10^−29^, t(29) = −47.5, paired sample t-test*, n = 30 APs, **Figure 4H**).

Altogether, these observations are in agreement with the expected filtering, which affects fast signals, such as the axonal APs more prominently.

Our model predicts that APs of the small MF bouton had lower peak and slower kinetics than APs measured from large MF terminals (Henrik Alle et al., 2009; Geiger & Jonas, 2000). These observed differences could derive either from biological variability among the subcellular compartments or from the larger deteriorating instrumental impact on the local membrane of the smaller axon. To discriminate between these possibilities, we investigated the *native* AP parameters predicted by the model. In this arrangement, we run the modeled biological structure with the reconstructed model conductances but the recording instruments was removed (**Figure 4G and H**). Thus, we predict how the native APs would look if the recording instrument was not present. We found that the native APs reached more depolarized peak potential (46.9 ± 0.09 mV, *p = 2.27*10^−48^, t(29) = 221.43, paired sample t-test*, n = 30 APs) and were significantly faster (half-width: 0.33 ± 0.002 ms, *p = 1.36*10^−33^, t(29) = −68.3, paired sample t-test;* dV/dt_max_: 1064.81 ± 5.1 V/s, *p = 3.75*10^−45^, t(29) = 171.47, paired sample t-test*, n = 30 APs) compared to the local spikes modeled with the experimental rig. The obtained native AP parameters were similar to those reported from large MF boutons, suggesting that the different AP shape observed in the small MF recording is primarily attributable to the presence of the measuring system.

Altogether, these results show that it is possible to obtain the native and local AP properties and the plausible underlying mechanisms using a complex instrumental model.

### Modeling the recording instruments accurately predicts signal distortions and native AP shapes

We used additional experiments to verify the reliability and plausibility of the model predictions. Specifically, we recorded APs with three different capacitance neutralization settings from the same axon and tested whether it alters the output of the model (CPN=6.5, 7 and 7.384 pF **Figure 5A**). Sub-optimal CPN conditions affects the recorded signal in several ways (**Supplementary Figure 4**). The simulations reproduced the recorded APs with different level of distortions when the CPN in the model was adjusted accordingly (**Figure 5A**). In addition, the predicted underlying sodium and potassium conductances were also very similar (g_Na_ density: 159.02 ± 1.73 mS/cm^2^, 151.7 ± 0.95 mS/cm^2^, 160.03 ± 1.47 mS/cm^2^, g_K_ density: 5.83 ± 0.05 mS/cm^2^, 5.41 ± 0.03 mS/cm^2^, 6.32 ± 0.09 mS/cm^2^; n = 30, n = 30 and n = 28 for CPN = 6.5 pF APs, CPN = 7 pF APs and CPN = 7.386 pF APs, respectively, **Figure 5B** and **Supplementary Figure 5**).

**Figure 5.**
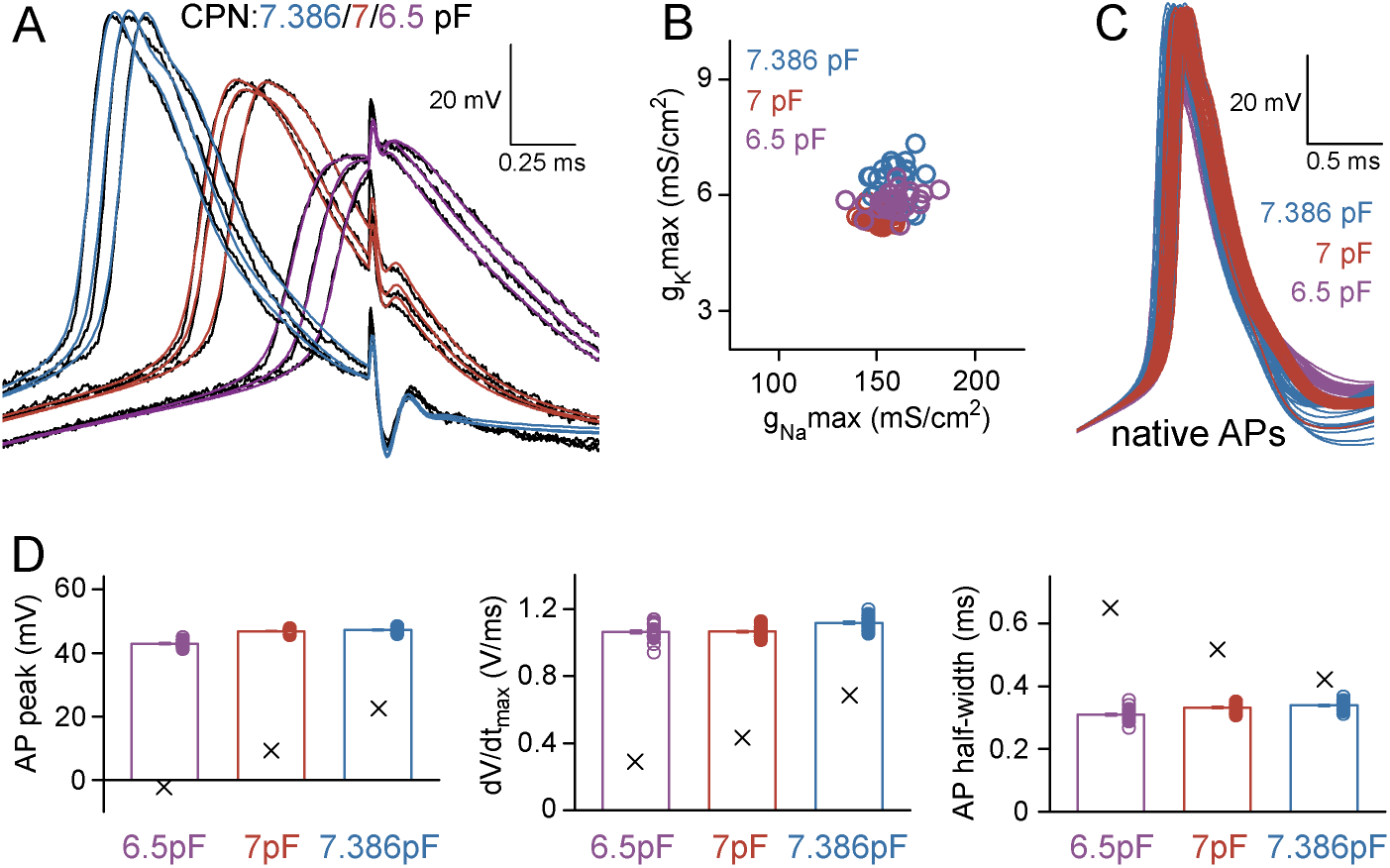
APs recorded and simulated from the same axon under different instrumental distortions predicted similar native spike shapes. (A) Representative target APs (black) and their best-fit representations in the complex instrument+axon model with standard CPN settings (red, 7 pF), with slightly reduced CPN (purple, 6.5 pF) or with the highest attainable CPN level (blue, 7.386 pF). (B) The best-fit gNa_max_ and gK_max_ were similar from APs with different CPN levels (n=30, 30 and 28 target APs). (C) The native AP waveforms retrieved from APs with different CPN levels were also similar. (D) Peak, maximal dV/dt and half-width of the native APs predicted based on recordings with three different CPN settings. Columns show the averages of the native APs, while X denotes the averages of the experimentally measured parameters (n=30,30 and 28 APs) which were distorted by the instruments.

Importantly, not only the predicted AP shapes matched but the native APs were also similar despite of the different levels of distortions in the three independent original recording conditions (absolute AP peak: 43.0 ± 0.17 mV, 46.9 ± 0.09 mV, 47.2 ± 0.14 mV, AP half-width: 0.31 ± 0.003 ms, 0.33 ± 0.002 ms, 0.34 ± 0.003 ms; n=30, n=30, n=28 for CPN=6.5 pF APs, CPN=7 pF APs and CPN=7.386 pF APs, respectively; **Figure 5C and D**). For an additional verification, we tested whether the model provides consistent predictions of AP propagation across conductance sets derived from different recording configurations. Specifically we simulated the natural AP conduction of distally evoked APs to the original recording site (423 μm away, **Figure 6A**) and measured the speed of propagation and the shape of the incoming APs. The incoming propagating APs were similar (absolute AP peak: 61.73 ± 0.13 mV, 64.22 ± 0.07 mV, 65.48 ± 0.08 mV, AP width at −10 mV: 0.34 ± 0.003 ms, 0.36 ± 0.002 ms, 0.33 ± 0.002 ms; n=30, n=30, n=28 for CPN=6.5 pF APs, CPN=7 pF APs and CPN=7.386 pF APs, respectively; **Figure 6**).

**Figure 6.**
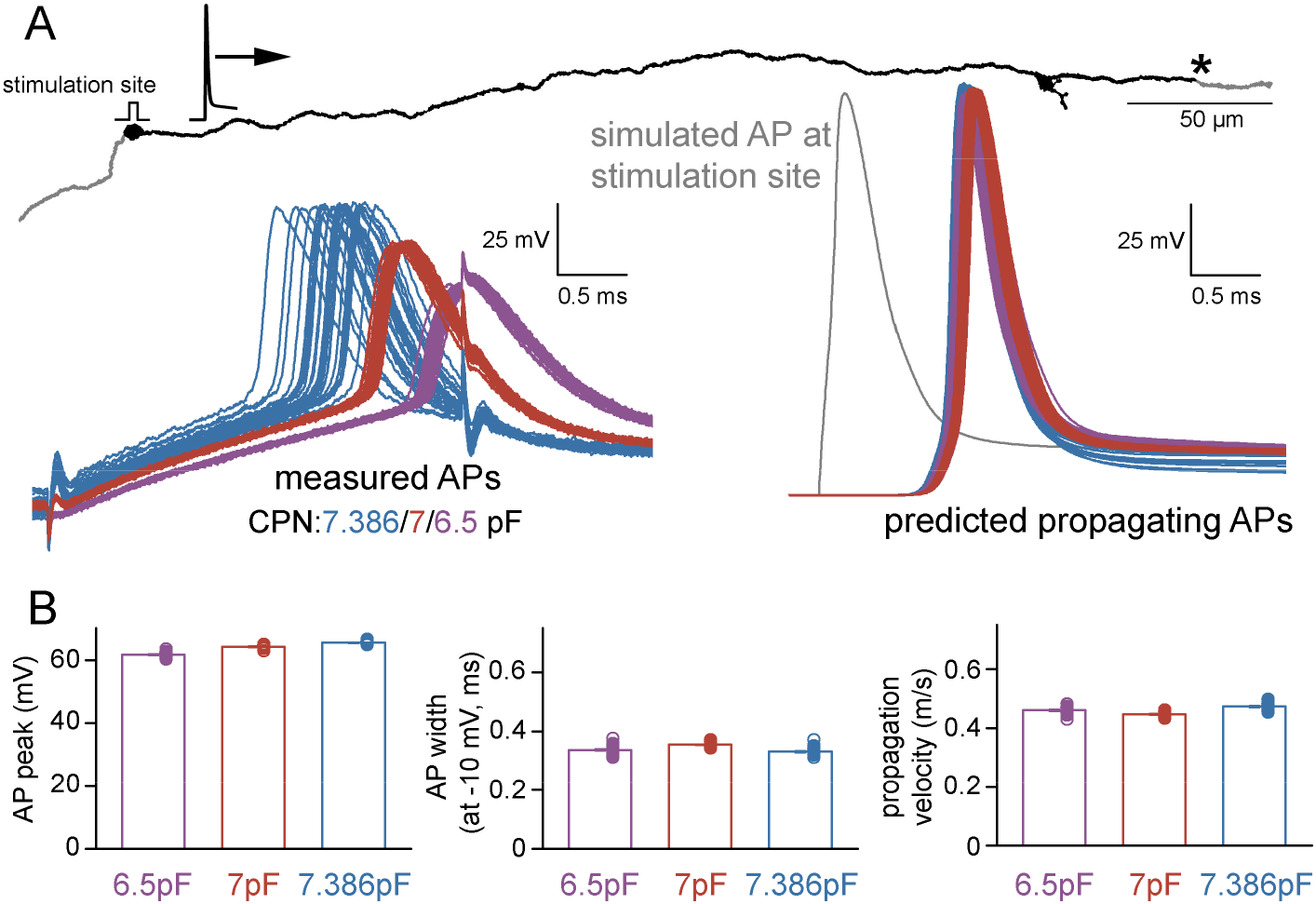
Characteristics of propagating native APs. (A) The reconstructed and simulated axonal structure with the position at which native AP parameters were captured (indicated with asterisk) after distal AP initiation (stimulation site) in individual models that were optimized for different recording conditions. Left, 88 original AP recordings that were used for the AP optimization in the models and recorded individually with three different CPN settings (n=30, 30 and 28 with 6.5, 7 and 7.368 pF CPN, respectively). The retrieved conductance sets were applied to the complete axon individually in all 88 cases. The right panel shows the 88 native, propagating simulated APs at the indicated distal point along the axon. The color is as for the measured APs on the left. A simulated AP at the stimulation site is shown in gray. (B) Peak, width at −10 mV and propagation velocity of the distally initiated propagating APs retrieved from 88 individual simulations. Note that conventional AP half-width measurements can not be applied for propagating APs because of the altered apparent threshold.

The prediction of AP propagation velocity is particularly sensitive to model parametrization as the speed of axonal spike propagation strongly depends both on the morphological properties of the axon and the specific passive and active mechanisms of the axonal membrane. Consistent with previous measurements in hippocampal MFs (Henrik Alle et al., 2009; Kress, Dowling, Meeks, & Mennerick, 2008), all model configuration predicted similar propagation speed (0.46 ± 0.002 m*s^−1^, 0.45 ± 0.001 m*s^−1^, 0.47 ± 0.002 m*s^−1^; **Figure 6B**).

Taken together, these results confirm that complete representation of the recording instruments in a model is sufficient for generating plausible native signals and underlying membrane mechanisms from signals that are distorted by the recording apparatus.

### Instrumental and structural parameters jointly define signal distortion in recordings from small neuronal structures

In addition to providing a useful tool for predicting and correcting instrumental distortions our simulations confirmed that complete elimination of the instrumental disturbance was not possible during recordings (Ritzau-Jost et al., 2021), since substantial difference persists between the measured and native APs, even when the capacitance compensation reached the maximally attainable level (see **Figure 5C**). The model raised the possibility that inadequate local signal generation also significantly contribute to the alterations in AP shapes in addition to filtering that affects recordings through patch pipettes. We refer to this effect as observer effect based on the analogy with the concept introduced in the field of physics for situations where the measurement inevitably changes the measured parameter. The observer effect can be quantified as the difference between local APs (signals in the structure when pipette is present) and native APs (signals without the presence of any instrument, **Figure 4G and H**). We quantified the relative contribution of filtering and observer effects to the total instrumental distortion in different experimental situations. Specifically, we varied pipette parameters, compensation settings and the size of the recorded cell (size was set to as Ccell = 1 pF and 10 pF, corresponding to the size of axons or small caliber dendrites and small cell bodies, respectively) in a reduced model which included only a single neuronal compartment with the instrumentation and compared the half-widths of APs (**Figure 7**). First, we isolated the filter effect by comparing simultaneous AP signals in the pipette and in the cell (measured versus local AP).

**Figure 7.**
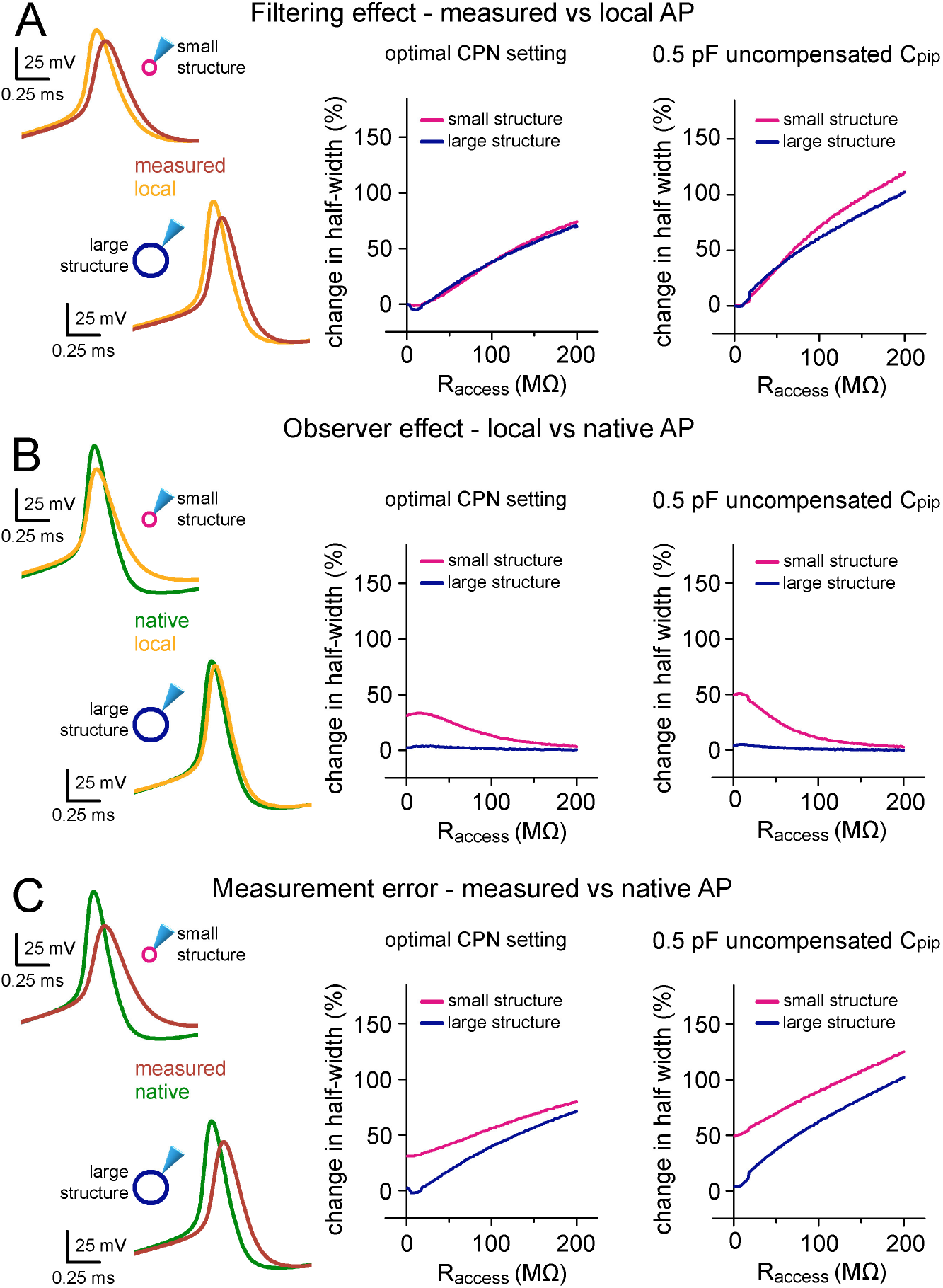
Instrumental and structural parameters cooperatively determine signal distortions in recordings from small neuronal structures. (A) On the left, the differences between the waveforms of the same APs within the pipette (measured) and in the two hypothetical cells (local) highlight the filtering effect of 70 MΩ R_access_. Only the size of the membrane surface were different in the two spherical structures, resulting in 1 and 10 pF biological capacitance, which correspond to small axonal and small somatic recordings. Graph in the middle summarizes the filtering effect quantified as the difference in AP half-width over a wide range of R_access_ in the small and large spherical cells. The right graph summarizes the filtering effect in simulations where suboptimal capacitance neutralization was applied (6.3 pF instead of 6.8 pF). (B) Using the same simulation environment as in panel A, the observer effect was quantified as the difference between the local AP and the native AP waveform. Thus, this data represents the isolated influence of the instrument on local signal generation. (C) To quantify the measurement error we demonstrate the difference between the measured AP and the native AP waveform in the same conditions as above. Thus, this is the sum of the filtering and observer effects.

In accordance with the general notion (Barbour, 2014), high R_access_ significantly filters fast voltage transients and allowed faithful measurement only in a confined R_access_ range (R_access_ that causes 10% AP signal widening: 47 MΩ for small cell and 41 MΩ in case of larger cell; **Figure 7A, left**). Furthermore, when uncompensated capacitance was added to the circuit by reducing the applied capacitance neutralization (−0.5 pF), the recording became more vulnerable and reliable recordings needed better R_access_ (R_access_ with 10% distortion: 20 MΩ for small cell and 17 MΩ in case of larger cell; **Figure 7A, right**).

Next, we isolated the observer effect by comparing the local and native AP shapes with or without the presence of the recording instrument (**Figure 7B**). The observer effect was always negligible in the case of the larger simulated biological structure as the capacitance added by the pipette was insignificant compared to the cellular parameters. In contrast, when the structure was smaller the structure had to discharge the remaining instrumental capacitance, which was in in this case in the range of Ccell, resulting in significant observer effect on the recorded AP shape (**Figure 7B**). Consistent with this hypothesis, observer effect was larger in simulations with additional uncompensated C_pip_ (mean change in AP half-width in the R_access_ range of 1-50 MΩ: 31.03±0.32 % vs. 40.54±1.14 %, optimal CPN vs. CPN with 0.5 pF uncompensated C_pip_, respectively; **Figure 7B**). Intriguingly, the observer effect showed a reversed dependence on the R_access_. It was the most pronounced when R_access_ was low. The larger R_access_ presumably isolates the residual instrumental capacitance from the cell and consequently weakens the instrumental impact.

Finally, by comparing the measured and native signals we assessed the overall measurement error. In case of the larger simulated structure, the overall measurement error is dominated by the filtering effect (**Figure 7C**). In contrast, when Ccell is small, the observer effect contributes significantly to the total error. Interestingly, the complementary changes in the two effects make these recordings less sensitive to changes of R_access_ and the error is similar when the R_access_ is low or high (**Figure 7C left**). The results confirm that residual uncompensated pipette capacitance further deteriorate the difference between native and measured AP signals (**Figure 7C left**). Similar conclusions can be drawn regarding other AP parameters as well (**Supplementary Figure 6**). Altogether, these model simulations demonstrated that measurement errors in patch-clamp recordings depend not purely on the pipette parameters but the size of the recorded structure itself has influence on the signal distortion.

Finally, to confirm the above findings on the error sources in a native biophysical structure we repeated the above tests in a reconstructed MF that was labeled during somatic recording that ensured more intact axonal arborization (**Figure 8A**). In this case the simulations ran with fixed instrumental parameters (6.74 pF C_tot_ and 60 MΩ R_access_, with 6.8 pF capacitance neutralization and 60 MΩ bridge balance correction; **Figure 8A, B and C**). We evoked APs along the axon and compared the pipette measured, local and native AP waveforms to determine the contribution of the observer and filtering effects to the instrumental signal distortion and their dependence on the local biophysical environment (**Figure 8A, B and C**). In agreement with the single compartment simulations (**Figure 7**), observer effect had a major contribution to the overall measurement error (change in AP half-width: 7.02 ± 1.02 %, 34.03 ± 4.59 % and 43.26 ± 4.51 % for pipette filtering, for observer effect and for total measurement error, respectively, n=941 locations; **Figure 8A, B and C**). As it is expected from the fixed instrumental parameters, pipette filtering was similar in small and large diameter sections of the axon. In contrast, the size of the observer effect was much larger in smaller axonal structures. Consequently, the measurement error was also the largest in the smallest diameter sections (**Figure 8B**). Importantly, minima in observer effect coincide with larger local capacitance, which originate from the large mossy fiber boutons in the CA3 area and branching points in the hilus. Indeed, the overall measurement error showed similar inverse correlation to local capacitance (R^2^=0.68, power) as the observer effect (R^2^=0.75, power) while the pipette filtering was structure-independent (**Figure 8D-F**). These findings confirm that measurement error critically depends on the local biophysical environment.

**Figure 8.**
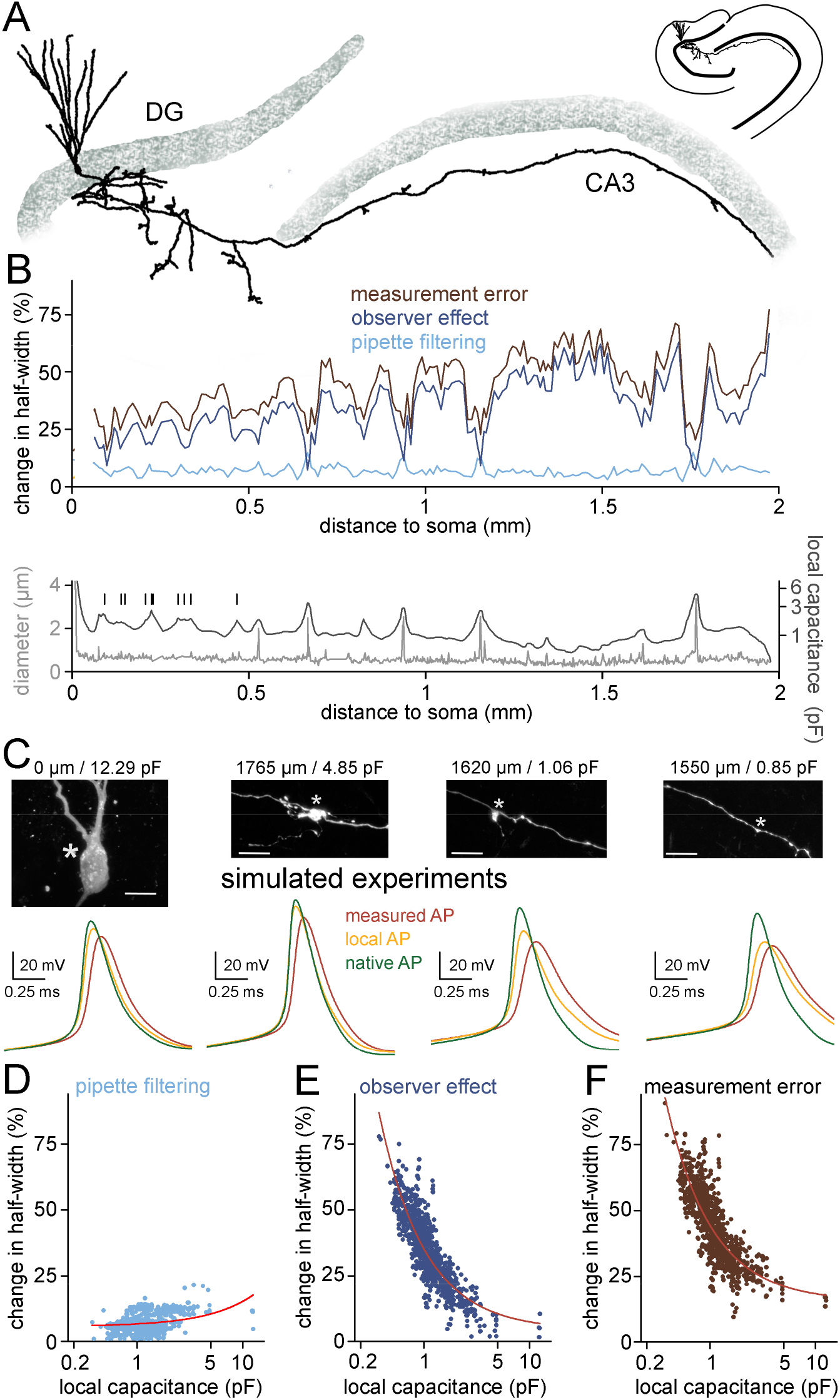
Measurement error along the axon is inhomogeneous and depends on the local biophysical environment. (A) Morphology of a somatically labeled complete granule cell that was used for simulating hypothetical recordings along its axon. (B) Top, observer- and filtering effects (dark blue and light blue, respectively) together with the overall measurement error (brown) in simulated axonal measurements plotted against the somatic distance of the recording position. Each value represent average of 5 individual measurements in each 10 μm segments. Bottom, axonal diameter (grey) and local capacitance (black) as a function of somatic distance. Vertical bars indicate the branch points of hilar axon collaterals. Notice that signals that derive from axonal segments that are closer to the soma are less affected by the observer effect due to the capacitive “load” of the somatic membrane. (C) Representative simulated recordings from the soma and different axonal sites and confocal images of the simulated recording site (same cell as in panel A). Corresponding somatic distances and local capacitances are indicated on the top. Asterisks mark the recording positions in the simulations. Scale bar=10 μm. (D) The filtering effect showed only weak correlation with the local axonal capacitance (n = 941 simulated recordings, R^2^ = 0.13, linear fit). Circles represent the filtering effect at independent measurement sites along the same axon. Note the logarithmic scale of the x-axis. (E) The observer effect showed significant correlation with local axonal capacitance (n = 941 simulated recordings, R^2^ = 0.75, fitted with power function). (F) Correlation between the total measurement error and the local axonal capacitance (n = 941 simulated recordings, R^2^ = 0.68, fitted with power function).

## Discussion

Here we developed a measurement-based, highly realistic model representation of the recording instrument that faithfully replicates actual patch-clamp recordings when combined with the detailed morphological reconstruction of the recorded structure. Simulation of the complete experimental condition allows for eliminating the instrumental distortions that are inevitably contaminates signals from small neuronal processes, such as axons. This realistic model also enabled us to determine the extent and sources of perturbations that contaminate electrophysiological signals. The results showed that physical parameters of the measuring instrument and the local biophysical properties of the recorded structure and their quantifiable interactions define the errors in voltage measurements. Consequently, signal deterioration potentially arise from the altered local signal generation instead of pipette filtering in small axonal recordings.

The core hypothesis behind our study is that realistic *in silico* representation of recording instruments together with the detailed morphology and biophysics of the recorded structure provide a better understanding of signal distortions present in direct voltage recordings and offers an applicable offline approach to predict native signals from distorted recordings. The combined simulation of the experimental conditions and the morphology was proved previously to be an ideal tool to describe technical limitations associated with VC measurements and to correct the recorded signals from those distortions (Beaulieu-Laroche & Harnett, 2018; Major et al., 1994; Schaefer, Helmstaedter, Sakmann, & Korngreen, 2003; Silver, Traynelis, & Cull-Candy, 1992; Spruston, Jaffe, Williams, & Johnston, 1993). However, these simulations have not included the complete recording instruments, which are known to impose significant distortions on the recorded signals, especially when the biological source is physically small, such as small axons. Other efforts corrected distortions of measured voltage signals by estimating the transfer functions of the specific recording instrument (Brette et al., 2008; Jayant et al., 2017; Magistretti, Mantegazza, de Curtis, & Wanke, 1998). Although such formulations are computationally efficient, they could not be applied to the variable individual contribution of circuit components (e.g. pipettes, C_pip_ and R_access_ compensatory elements), which imposes different signal distortions, such as filtering or instrumental capacitive load. Here, we implemented the pipette and all amplifier elements as individual circuits as they were actually built in the complete system and showed that the idealized instrument is sufficient to replicate the behavior of the complete measuring instrument and its compensatory capabilities (**Figures 1, 2 and 3**). Thus, we assume that implementing these components will allow adapting the model to other amplifier types. One of the unexpected findings of our study is the uneven capacitance distribution of recording pipettes (Benndorf, 1995). Majority of the total C_pip_ originate from the first two and a half mm from the tip (**Figure 3**). The gradual increase of relative wall thickness from the tip explains the larger contribution of the tip. Although the relative wall thickness was found to be constant previously for borosilicate glass capillaries (Benndorf, 1995) we observed decreased relative wall thickness toward the tip that explains the larger capacitive contribution of the tip.

To outline a feasible application, we utilized the complete instrumental model to correct the shape of APs directly recorded from a small axon terminal of a hippocampal mossy fiber. The size of this bouton (largest diameter: 0.7 μm) was in the range of typical cortical axon terminals and was much smaller than the famous large mossy fiber terminals. The complete instrumental model in combination with the realistic axonal morphology and biophysics faithfully replicated the measured voltage signals (**Figures 4, 5 and 6**), including the signal artefacts and distorted fast APs. The simulation resulted in ionic currents and conductances that matched with previous results obtained with direct MF recordings from large boutons (**Figure 4E and F**) (H. Alle et al., 2011; Henrik Alle et al., 2009; Geiger & Jonas, 2000). Consequently, the retrieved native axonal APs were brief events with large amplitude (**Figures 4, 5 and 6**) whose shape closely resembled to the spike waveforms reported for the large axon terminals of the same axon (Henrik Alle et al., 2009; Geiger & Jonas, 2000). The simple Hodgkin-Huxley type conductance models with six free parameters were sufficient to restore the experimentally recorded MF AP waveforms. Although APs in the distal MFs can be described with a similar standard Hodgkin-Huxley type gating (Engel & Jonas, 2005; Ohura & Kamiya, 2018), AP simulation in other neuron types or in other subcellular elements required more detailed kinetic schemes (Stefan Hallermann et al., 2012; Ritzau-Jost et al., 2014; Ritzau-Jost et al., 2021; Schmidt-Hieber & Bischofberger, 2010). Therefore, implementation of more elaborated conductance models and inclusion of additional conductances can further improve our simulations and adapt to multiple activity regimes, such as the plasticity of AP shapes during sustained activity (Geiger & Jonas, 2000).

The complex model allowed us to dissect the sources of errors that contaminate recordings from small biological structures. As previously described for small, electrotonically compact neurons (D’Angelo, De Filippi, Rossi, & Taglietti, 1995; Goodman, Hall, Avery, & Lockery, 1998; Rohrbough & Broadie, 2002), we found that even with careful capacitance neutralization, the capacitive load added by the recording instrument substantially altered the intrinsic electrical behavior of the axon (**Figures 4 and 5**). Using a simplified neuronal representation, we confirmed with that instrumental capacitance effectively attenuates the action potentials in small neuronal structures (**Figure 7**) emphasizing again that C_pip_ reduction can substantially improve the accuracy of the measured voltage signal (Dudel, Hallermann, & Heckmann, 2000; Levis & Rae, 1993; Ogden & Stanfield, 1994; Ritzau-Jost et al., 2021; Sakmann & Neher, 1983). Interestingly, we have shown that increase in the R_access_ reduced the instrumental impact on the cellular electrogenesis (probably due to effective electrical isolation of the neuronal structure from the recording pipette) suggesting that high impedance recording can have also advantages when experimental subject is small. The high impedance recordings may reduce the complexity of *post hoc* AP reconstitution as in this case the predicted AP shape depends only on the pipette filtering (Jayant et al., 2017). One of the major findings of our study is that instrumental and cellular factors define the accuracy of CC experiments not in isolation from each other, but their interaction is equally important. This was the most apparent when we examined the cellular contribution of the overall measurement error along a reconstructed MF which forms varicosities with different diameter. Thus, the differently sized axon segments provides different local biophysical conditions (**Figure 8**). In this arrangement, which is characteristic to all axons at a various degree (i.e. the size of the terminals and axonal shaft varies), the observed inverse relationship between local capacitance and the measurement error highlight the importance of structural details on direct axonal recordings.

A more general consideration is that the target-size-dependent effects are not specific to the axonal recordings. Experiments that target small cellular structures, whose electrical parameters are comparable to the capacitance introduced by the measuring instrument, are potentially subject to the distortions quantified in our study. The target-dependent measurement errors are not specific to the action potential firing either. In fact, the typically high conductance densities in axons (Hu & Jonas, 2014; Ritzau-Jost et al., 2014) can partly compensate for the observer effect. Depending on the local biophysical environment, the recording instruments can induce substantial observer error in dendritic membrane potential as well.

## Methods

Experimental procedures were made in accordance with the ethical guidelines of the Institute of Experimental Medicine Protection of Research Subjects Committee (MÁB-7/2016, PE/EA/48-2/2020).

### Constraining the amplifier model

#### The model cell

Electrical components of the customized model cell (test#3 circuit, modified type 1U, Molecular Devices) were connected through conductive metal slots taken from a circuit breadboard allowing the change of circuit components without the need for soldering that would introduce variable stray capacitance (**Supplementary Figure 1**). We used non-inductive, low-noise resistors: resistors either originally present in the 1U model cell, 10MΩ and 500 MΩ, or Ohmite SLIM-MOX10203 series, 20-100MΩ. Stray capacitance of each elements of the test circuits (including the resistors and their slots) was characterized in VC mode by measuring the capacitive load associated with the introduction of the given circuit element. All capacitors were considered ideal, that is, without any resistive component.

#### Boosting unit

We implemented a simplified boosting unit (Sigworth, 1995) in which capacitors were fixed (100 pF and 120 pF) while resistors were directly fitted to reproduce the capacitive current response in the test#1 configuration. Late phase of the response profile (starting 24 μs after the stimulus onset) has particular importance because artefacts in that temporal domain can contaminate measured biological signals, therefore, this part of the signal was heavily weighted (850x) during the adjustment of the boosting unit.

#### VC capacitance compensation

Both CPf and CPs circuits were designed as described previously (Sigworth, 1995).

#### Stray capacitance of the CC circuit

To optimize the circuitry of CC model, we performed measurements with test#3 circuit, where pipette parameters (10MΩ, 2.8 pF) and the size of the cell-equivalent resistor (500 MΩ) were fixed, only cellular capacitance varied from 0.75 pF to 46.7 pF. We applied short current stimuli (−50 to −200 pA, 3 ms) to elicit voltage responses with the maximal attainable capacitance neutralization or without CPN. Traces recorded in the absence of CPN allowed us to characterize the total capacitive load of the CC circuitry. The model most accurately reproduced the real voltage responses when a 0.76 pF stray capacitance was added at the amplifier input node (C_CC_ on **Figure 1**).

#### Capacitance neutralization (CPN)

We implemented CPN in two steps. First, we added an idealized positive feedback loop, where the compensation through the C_inj_ can be modulated with the gain of an operational amplifier. This simple circuit representation was sufficient to reproduce the neutralizing capability of the real amplifier, that is, equal CPN settings resulted in the same level of compensation in the model as in the real experiments. Next, we reproduced the characteristic CPN-related stimulus artefacts that are present in typical current clamp measurements. We placed a resistor (R_CPN_) and an inductor (L_CPN_) to the CPN path (**Figure 1**) and tuned their parameters by direct fitting of the model voltage responses to the experimental data (examples are shown on **Figure 2D**).

#### Bridge balance compensation (BB)

To recreate the BB circuit, we created a reference signal from the command equivalent to the voltage drop caused by 1 MΩ load resistance (**Figure 1**). Scaled version of that reference signal was then subtracted from the pipette voltage for the correction.

#### Bessel filters

We added filters to the amplifier model to match of our actual biological recordings. We added an active linear filter consisting of two cascaded Sallen-Key filter stages (**Figure 1**). Filter parameters were set according to an available filter design tool (https://www.analog.com/designtools/en/filterwizard/) to produce output with four pole low-pass Bessel filter characteristics. All recordings were made with bypassed filtering mode and we applied 100 kHz lowpass filter throughout the simulations.

### Pipette parameter measurements

We used typical borosilicate glass capillaries (BF150-75-10, Sutter Instruments, Novato, CA) to fabricate pipettes that are suitable for recordings from small axons. To implement these high impedance pipettes into the model we measured their actual parameters. First, we assessed the actual pipette capacitance as a function of the tip distance by dipping known part of the recording pipettes into the recording solution. Pipette position was measured by the x axis values of the motorized micromanipulator (SM5 controller with Mini unit, Luigs und Neumann, Ratingen, Germany). First, we recorded the total capacitance of the instrumentation in open circuit VC mode when the pipette was out of the solution. Then, we gently moved the pipette to a position where the tip intermittently reached the surface of the fluid characterized by the appearance of short conductive periods in the recorded VC signal. Then, the pipette was pushed forward to the solution to reach a position where the conductive state became stable (typically <5 μm forward movement). Starting from this tip position point, we systematically increased the length of the dipped part and quantified the capacitance in VC by integrating the first 50 μs of the transient response to a −20 mV voltage step. Integrated area was divided by the voltage step amplitude to convert the electric charge to capacitance.

To measure distribution of resistance along the pipettes, we broke off a known length of the pipette and measured the resistance of the remaining part in VC mode. First, we moved the pipette tip to a defined position under the objective and recorded its resistance. After withdrawal of the pipette, we broke the tip by gently touching it with a piece of lens cleaning tissue. The newly formed pipette tip was then positioned back to the reference position on the image. We determined the length of the broken pipette parts by reading the difference of the positioning motor. Resistance was measured in VC mode using −2 to −20 mV steps. We repeated the breaking process for each pipettes several times (1x-8x, using 22 pipettes in total).

We visualized the outer/inner diameter ratio (R_OI_) of the recording pipettes using a grinding system that gradually cut them in half. This measurement allowed to avoid the optical distortions caused by the lens effect of the cylindrical glass. The recording pipettes (the first 8-12 mm from the tip) were embedded into epoxy resin on a microscope slide. Pipettes were longitudinally grinded with a coarse-grained aluminium-oxide abrasive disc (grit size=600) until we reached the surface of the pipettes. Then, the grinding was occasionally interrupted to check the surface and cross-section of the pipette under a transmitted light microscope (Leica DM2500 microscope, 5X-100X magnifications). As the plane of the pipette tip was approached, we switched to fine-grained abrasive discs (grit size=6000). The images obtained from different cross section levels were then used to measure the pipette diameters (Adobe Photoshop 5.0). The diameter data were included only from those focal planes for distinct segments of the pipette tips, where the outer diameter was the largest and the relative wall thickness was the smallest. We assumed infinite pipette wall resistivity in the model.

### Slice preparation and electrophysiology

Hippocampal slice was prepared from a 29 days old Wistar rat as previously described (Brunner & Szabadics, 2016). In brief, the animal was deeply anaesthetized with isoflurane. After decapitation, the 350 μm thick slice was cut with Leica VT1200S vibratome in ice-cold cutting solution (85 mM NaCl, 75 mM sucrose, 2.5 mM KCl, 25 mM glucose, 1.25 mM NaH_2_PO_4_, 4 mM MgCl_2_, 0.5 mM CaCl_2_, and 24 mM NaHCO_3_) in an orientation optimized to preserve the mossy fibre tract in the hippocampal CA3 area (Bischofberger, Engel, Li, Geiger, & Jonas, 2006). The slice was incubated at 32 °C for 60 minutes after sectioning, and was stored at room temperature until the experiment. The recording solution was composed of 126 mM NaCl, 2.5 mM KCl, 26 mM NaHCO_3_, 2 mM CaCl_2_, 2 mM MgCl_2_, 1.25 mM NaH_2_PO_4_, and 10 mM glucose (equilibrated with 95% O_2_ and 5% CO_2_ gas mixture). The pipette was filled with an intracellular solution containing 90 mM K-gluconate, 43.5 mM KCl, 1.8 mM NaCl, 1.7 mM MgCl_2_, 0.05 mM EGTA, 10 mM HEPES, 2 mM Mg-ATP, 0.4 mM Na2-GTP, 10 mM phosphocreatine, 8 mM biocytin and 20 μM Alexa Fluor 594 hydrazide (pH=7.25). The recordings were performed at 35 °C. For patching, axon was visualized with an upright IR-DIC microscope (Eclipse FN-1; Nikon) equipped with high-numeric aperture objective (Nikon 1.1 NA, Apo LWD 25 W), oil condenser (Nikon D-CUO DIC Oil Condenser, 1.4 NA) and a sCMOS camera (Andor Zyla 5.5). After the recordings the Alexa Fluor 594-labelled structure was visualized in situ using a confocal system (Nikon Eclipse C1 Plus). Then, the slice was fixed for further morphological experiments (see below).

To record from the individual small MF terminal, we searched for visually identifiable axonal structures under the guidance of the DIC optics in the *stratum lucidum* of the CA3 area, whose size was smaller than those of the typical large mossy fiber boutons. The patch pipette was pulled from borosilicate glass capillary (inner diameter: 0.75 mm, outer diameter: 1.5 mm, Sutter Inc.). After the seal formation, we manually carefully compensated the C_pip_ in VC mode. Compensated capacitive responses to −20 mV step command were then recorded and the average of 164 sweeps served as target for tuning the C_pip_ in the model. After establishing the whole-cell configuration by applying sudden negative pressure, we switched to CC mode to record the passive and active membrane responses from the axon with different CPN compensation. All recordings were collected with a MultiClamp 700B amplifier (Molecular Devices) without filtering (filter bypassed) and digitized with Digidata1440 A/D board (Molecular Devices) at 250 kHz sampling rate using the pClamp10 software package (Molecular Devices). At the end of the experiment, we collected preliminary morphological data by imaging the Alexa Fluor 594 signal with the confocal system. The obtained z stack image was then used to confirm the MF identity (evidenced by the presence of a large MF terminal along the recorded axon, **Figure 4B**) and for the documentation of the recording position.

### 3D morphological reconstruction

After the recording, slice was fixed overnight at 4°C in 0.1 mM phosphate buffer containing 2% PFA and 0.1% picric acid. The slice was then re-sectioned (to 60 μm) and the sections were incubated overnight with Alexa Fluor 594-conjugated streptavidin in 0.5% Triton X-100 and 2% normal horse serum to reveal the biocytin signal. To investigate the detailed morphology, the recorded axon was imaged with a confocal system (60X objective, Plan Apo VC, NA=1.45, Nikon C2, x-y pixel size:0.08-0.1 μm, z-step: 0.1-0.15 μm). High resolution reconstructions were done automatically by the Vaa3D software (Peng, Bria, Zhou, Iannello, & Long, 2014; Peng, Ruan, Long, Simpson, & Myers, 2010; Peng, Tang, et al., 2014).

### Implementation of the seal in the model

The seal was represented as a single resistor (R_seal_) connected in parallel to the cell. To estimate the R_seal_ in the axonal recording, first we calculated the ratio (R_ratio_) between the cellular input resistance (R_cell_) and the R_seal_ based on the voltage shift produced by the shunt conductance of seal (Perkins, 2006):

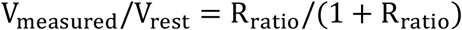

where V_measured_ is the recorded voltage, V_rest_ is the native resting membrane potential. R_seal_ and R_cell_ were then calculated from the measured input resistance (R_in,measured_) and from the R_ratio_ using the equations:

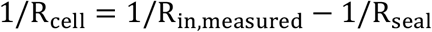

where:

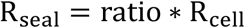

The apparent resting membrane potential of the recorded MF axon was −75.2 mV, close to the resting membrane potential previously reported for the somata and axons of granule cells (~ −80 mV (Brunner & Szabadics, 2016; Martinello, Giacalone, Migliore, Brown, & Shah, 2019; Ruiz, Campanac, Scott, Rusakov, & Kullmann, 2010; Schmidt-Hieber, Jonas, & Bischofberger, 2004; Staley, Otis, & Mody, 1992)). This moderate shift, caused by the leak through the seal, suggests orders of magnitude larger R_seal_ compared to the axonal input resistance. Indeed, we calculated 65.64 GΩ R_seal_ and 3.94 GΩ R_cell_ for the recording assuming −80 mV resting membrane potential. Accordingly, we applied 65.64 GΩ R_seal_ in the AP reconstitution model (**Figures 4, 5 and 6**), while R_seal_ was set to 50 GΩ in all other simulations. To compensate for the seal-induced depolarizing voltage shift, baseline membrane potential in the model was adjusted by constant current injection to match those of the experimental data.

### General simulation parameters

Cellular membrane parameters were assumed to be spatially uniform, unless stated otherwise. Potassium and sodium equilibrium potentials were set to −77 mV and +70 mV, respectively and we assumed −80 mV reversal potential for the leak conductance in active models. Simulations ran with 4 μs and 0.5 μs temporal resolution, in CC mode and in VC mode, respectively.

### Fitting procedures

We fitted the model responses to the experimental data to minimize the sum of squared error between them using the Brent’s PRAXIS optimization algorithm embedded in NEURON. For C_pip_ estimation, a 5 ms long trace was considered with 2.5 ms baseline before the stimulus. For passive parameter estimations, we weighted the voltage response evoked by short current stimulus (25.8 ms from the stimulus onset, 3X) in order to equalize the contribution of short-and long pulse responses to the total error. For AP reconstruction, fitting interval started with 2.6 ms long baseline period before the stimulus onset. The actual APs were weighted eight-fold, starting 0.5 ms before the peak. The optimization ended 1.5 ms following the AP peak to avoid the contamination of the parameter estimation with afterdepolarization related mechanisms not implicated in the model (Martinello et al., 2019; Ohura & Kamiya, 2018).

We used sodium and potassium conductance mechanisms with canonical Hodgkin-Huxley gating scheme, in which not only the density can be freely adjusted but also the kinetics and the voltage dependence of the two types of conductance. For this, we modified the built-in Hodgkin-Huxley mechanisms of NEURON by introducing rate-scaling factors (scNa and scK) to adjust the speed of model channel operation. The originally implemented temperature scaling was removed from the mechanisms. In addition, we used global voltage shifts (vsNa and vsK) to modulate the voltage dependence of the channels. Altogether, free parameters (gmaxNa, gmaxK, scNa, scK, vsNa, vsK) were constrained to obtain an ideal model of the AP waveform measured in our various experimental settings. Because the R_access_ can change during the recordings, this parameter was set individually for each target trace. Optimization of the individual AP fits was initiated from four parameter sets that allows exploration substantial part of the parameter space (**Supplementary Figure 3A**). The parallelized codes for AP reconstitution were run on the Comet supercomputer through the Neuroscience Gateway portal (Sivagnanam et al., 2013). To assess the quality of individual fits, their mean squared error was normalized to the baseline variance of the actual target trace. Of the results of the four parallel optimizations, the solution where the normalized error was the smallest was accepted as the best fit. Fit was rejected when the normalized error was larger than 10. Fitting of 2 of the original 90 target APs did not resulted in solutions, which met this criterion. These APs were excluded from the analysis (both were obtained with 7.386 pF CPN).

### Assessments of the reliability of optimizations

*In silico* reconstruction of the complete experiment consisted of subsequent optimization steps (Supplementray Figure 2A). First, we used VC data to set actual pipette parameters (1). After adding the model instrumentation and the detailed morphology of the recorded structure to the model, we tuned the passive cellular parameters together with the R_access_ (2). Next, we equipped the model structure with active sodium and potassium conductances and adjusted their properties to match model responses the experimentally recorded AP waveform (3). Finally, having established the appropriate conductance sets, we obtained the native behavior of the axon (4). The reliability of the fitting procedures were tested with artificial traces with Gaussian noise.

To assess the accuracy of C_tot_ estimation, we generated target traces by simulating on-cell VC experiments. The model always recovered correct C_tot_ value (proportion of fitting estimations within 10% error to the correct value: 100 %, n=990 optimizations, **Supplementary Figure 2A and B**) irrespective to the C_tot_ (5.8-13.7 pF) and noise level (SD=0-35.3 pA) of the targets, which confirms the high sensitivity of VC based C_pip_ estimation.

To test the reliability of cellular passive parameter and R_access_ prediction, we created a hypothetical axon with biologically plausible diameter distribution (log-normal distribution with mean of 0.6 μm and variance of 0.4 μm^2^). We attached the model axon to the pipette and generated hypothetical CC measurements with long and short current stimuli (C_m_ range: 0.5 - 1.5 uF/cm^2^, R_m_ range: 10 - 100 kΩ/cm^2^, R_i_ range: 50 - 250 Ω*cm, R_access_: 50 - 400 MΩ, noise SD = 1.73 mV). We fitted these synthetic targets from a single initiation parameter set (C_m_: 1 pF/cm^2^, R_m_: 50 kΩ/cm^2^, R_i_: 150 Ω*cm, R_access_: 150 MΩ). 75 % of optimizations resulted in acceptable results, that is, the fit error was less than 10-times the baseline variance of its target. In those successfully fitted cases the predicted parameters were close to their predefined values (proportion of fitting estimations within 10% error to the correct value: C_m_: 90.83 %, R_m_: 97.66 %, R_i_: 81.88 %, R_access_: 82.3 % n= 469 optimizations, **Supplementary Figure 2C**).

We also verified the reliability of the prediction of native AP shapes in independent simulations (**Supplementary Figure 2D and E**). For this we generated diverse AP waveforms using independent, sophisticated sodium and potassium conductance mechanisms (using 8-state kinetic schemes obtained from (Stefan Hallermann et al., 2012; Schmidt-Hieber & Bischofberger, 2010) in a single compartment neuron (R_access_=100 MΩ, C_pip_=6.74 pF, BB compensation=100 MΩ, CPN compensation= 6.85 pF, R_seal_=50 GΩ, noise SD: 0.45 mV). Our standard fitting routine reliably retrieved the shapes of APs.

### Systematic errors in model predictions

We also examined the potential impact of systematic error sources on model predictions. Such potential error can originate from inaccuracies of the morphological reconstruction. Post hoc anatomical processing can result in considerable tissue shrinkage. Additionally, diameter of thin axonal processes is close to the resolution limit of light microscopy. To evaluate the potential effects of inaccurate anatomical representation of the recorded axon, we artificially altered the reconstructed morphologies and tested how the recovered native APs were affected. The axonal diameters were either homogeneously increased by 160 nm or reduced by 180 nm (axons were not allowed to shrink below 100 nm). As expected, recovered passive parameters and active conductance densities scaled with the diameter to compensate for the altered morphological dimensions (**Table 1**). However, as the model re-adjusted the local electrical environment by scaling the membrane properties, the predicted intra-axonal and native AP waveforms remained remarkably similar in spite of the large changes in the morphology and passive membrane parameter.

**Table 1.**
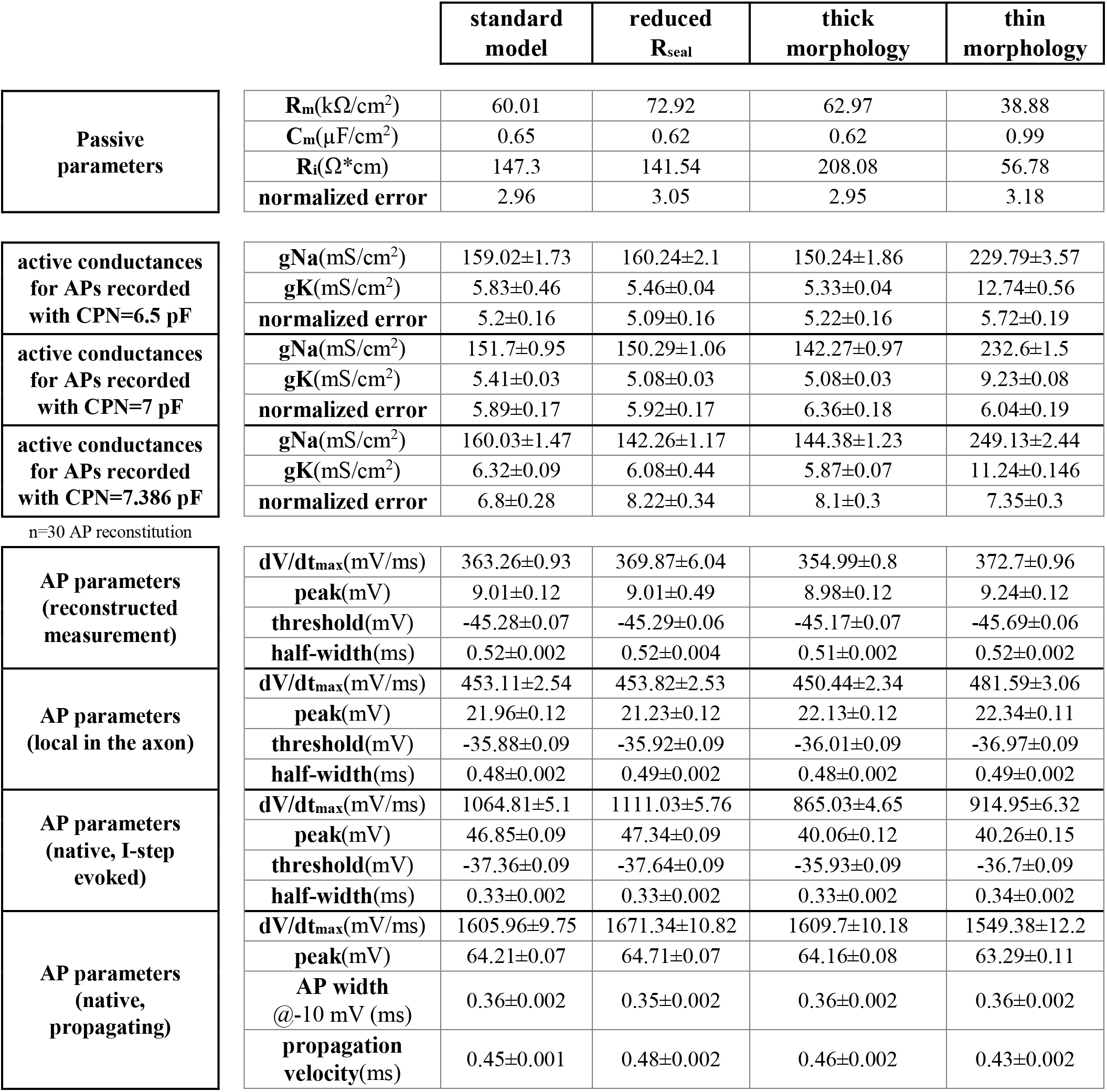
Summary of parameters obtained from different model configurations

*R_seal_* provides an additional error source because its calculation depends on the native resting membrane potential, which can not be assessed directly (i.e. measurements start in cell attached mode). Therefore, we defined the theoretical lower limit for the R_seal_ in the direct axonal recording; when native resting membrane potential equals with the potassium equilibrium potential (−93 mV with the solutions used in the experiment) R_seal_ would be only 20.58 GΩ. Re-optimization of the model using this small R_seal_ caused only minor alterations in the predicted conductances and AP parameters (reduced R_seal_ model, **Table 1**). Altogether, these control simulations suggest that potential systematic errors have marginal impact on the primary results of the manuscript.

### Simulations to explore the correlations between signal distortions and recording conditions (Figure 7)

We used a single compartment with fixed 10 μm^2^ surface area to represent neuronal structure in these simulations. Electrical behavior of small and larger cells was set by scaling the active and passive biophysical parameters of the compartment (for small cell: C_m_=10 μF/cm^2^, R_m_=2 kΩ/cm^2^, g_Na_ density=1.5 S/cm^2^, g_K_ density=0.4 S/cm^2^; for larger cell: C_m_=100 μF/cm^2^, R_m_=0.2 kΩ/cm^2^, g_Na_ density=15 S/cm^2^, g_K_ density=4 S/cm^2^). To keep R_seal_/R_cell_ ratio constant between the two conditions, R_seal_ was set to 50 GΩ and 5 GΩ, for small and larger cell, respectively. The pipette capacitance was fixed to 6.74 pF while R_access_ was systematically varied in the range of 1-200 MΩ. To mimic optimal recording configuration, the applied CPN settings (6.8 pF) closely matched with C_pip_. APs were elicited with 3 ms long current stimuli (30 pA and 160 pA for small and larger cell, respectively).

### Simulations to explore the correlations between signal distortions and axonal morphology (Figure 8)

The detailed morphology of a somatically labeled granule cell was imported to the NEURON. Dendritic spines were implemented by scaling two-fold the C_m_ and the leak conductance in dendrites. Passive parameters (Ri:150 Ω*cm, C_m_:1 μF/cm^2^, R_m_:50 kΩ/cm^2^) were constant otherwise. Active conductances (g_Na_ density=300 mS/cm^2^, g_K_ density=15 mS/cm^2^) were homogenously distributed along the cell, except the initial part of the axon where we applied higher channel densities (g_Na_ density=1200 mS/cm^2^, g_K_ density=75 mS/cm^2^) with left-shifted activation and inactivation (10 mV in the hyperpolarized direction). To explore the distortions in axonal AP recordings, position of recording electrode (C_tot_=6.74 pF, R_access_=60 MΩ, R_seal_=50 GΩ, applied CPN=6.8 pF) was systematically changed along the main axon (n=942 recording positions). APs were evoked by 3 ms long current stimuli at each recording position. Amplitude of the injected current was automatically adjusted to evoke APs ~1.5 ms (1.33 ± 0.01 ms delay, n=942 APs) after stimulus onset. Local capacitance was determined for each recording position using idealized VC simulations in the absence of experimental instrumentation (built-in SEClamp mechanism with 10 MΩ series resistance). For quantification, we integrated the first 100 μs of the transient capacitive response to −20 mV voltage step. The resulted charge was divided by the voltage step amplitude.

### Data analysis and statistics

AP threshold was determined as membrane voltage where depolarization rate exceeded 20 mV/ms. AP amplitude was calculated as the voltage difference between the absolute peak potential and the threshold. AP half-width was defined as the spike duration at half of its amplitude. AP conduction velocity was determined by measuring the temporal difference between AP peaks at the initiation and at the recording site.

Data were analyzed using pClamp (Molecular Devices), OriginPro (OriginLab) and Excel (Microsoft) Photoshop (Adobe) and custom written NEURON or Python scripts. The relevant NEURON codes are available on GitHub (https://github.com/brunnerjanos/amplifier-model). Voltage values are presented without correction for the liquid junction potential. Normality of the data was assessed with Shapiro-Wilks test. Population data are presented as mean ± s.e.m.

## Acknowledgements

This work was carried out with the full support of János Szabadics’ laboratory. We are grateful to János Szabadics for his continuous support, advices on the project and constructive comments on the manuscript. This work was funded by ERC-CoG 772452 (nanoAXON) grant to János Szabadics and the János Bolyai Research Fellowship of the Hungarian Academy of Sciences (to JB). We are thankful for the computational resources provided by the Neuroscience Gateway. We thank László Barna, the Nikon Microscopy Center at the Institute of Experimental Medicine, Nikon Europe B.V., Nikon Austria GmbH, and Auro-Science Consulting Ltd, for kindly providing microscopy support and Dóra Kókay and Andrea Szabó for technical assistance.

---

**Supplementary Figure 1.**
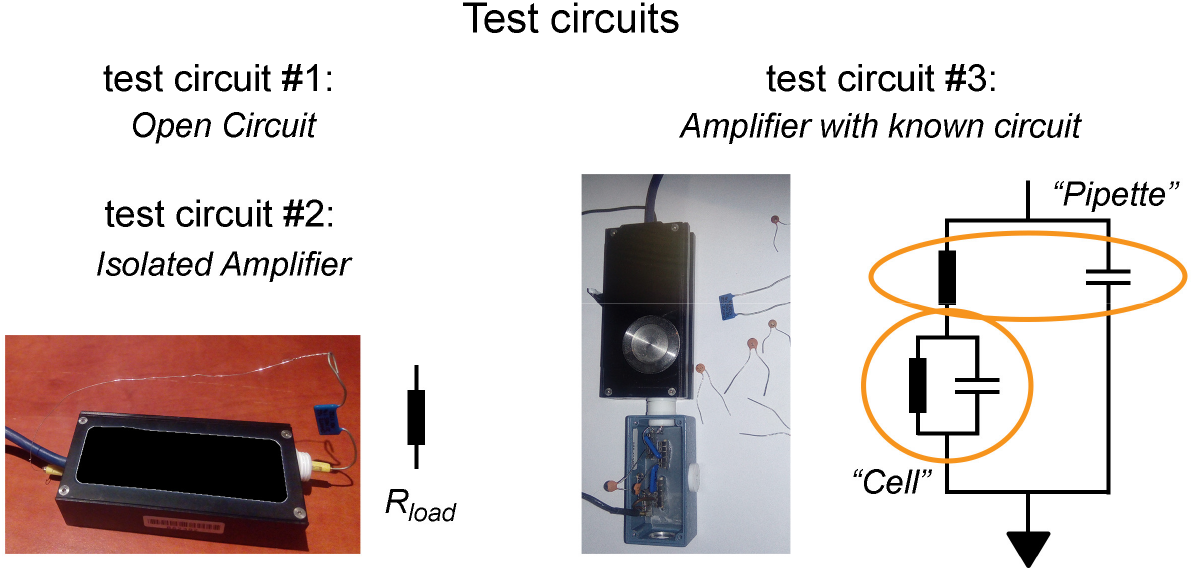
Supporting data for Figures 1 and 2. Test circuits used for the characterization of the circuit components.

**Figure S2.**
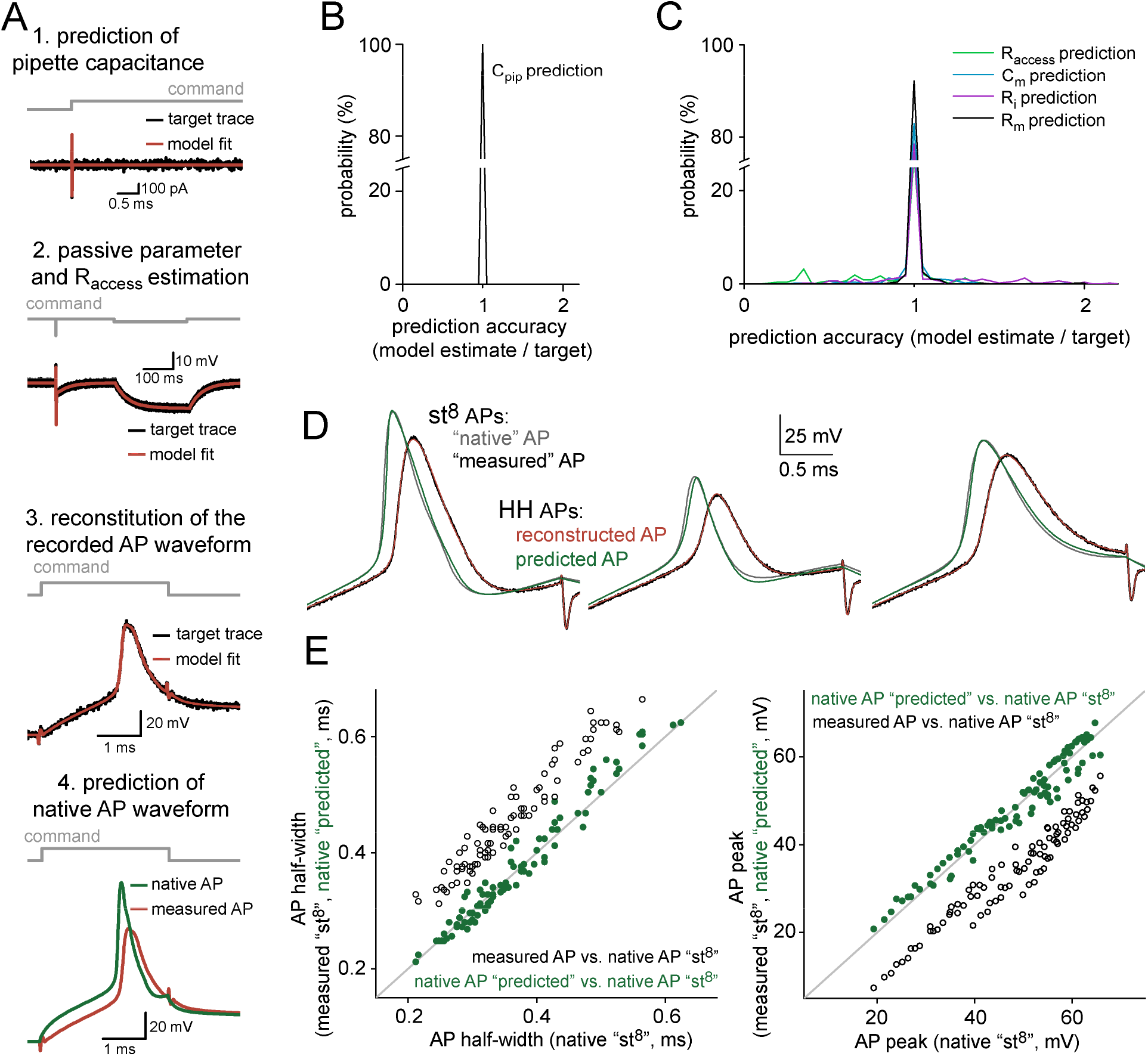
Supporting data for Figure 4. Parameter extraction from recorded VC and CC data. (A) Outline of the consecutive optimization steps that were used to retrieve the instrumental parameters. (B) Distribution of the relative error in C_pip_ estimations. (C) Distribution of the relative errors present in the R_access_ and passive parameter estimations. (D) Overlay of representative APs (native and measured, gray and black traces, respectively) generated using 8-state active conductance models and corresponding best-fit APs generated with the standard conductances used in our final model (reconstructed and predicted, red and green traces, respectively). (E) Comparison of measured AP half-width and peak values (black) and the corresponding predicted native parameters (green) with the original value (n=90 simulated experiments). The line indicates equality.

**Supplementary Figure 3.**
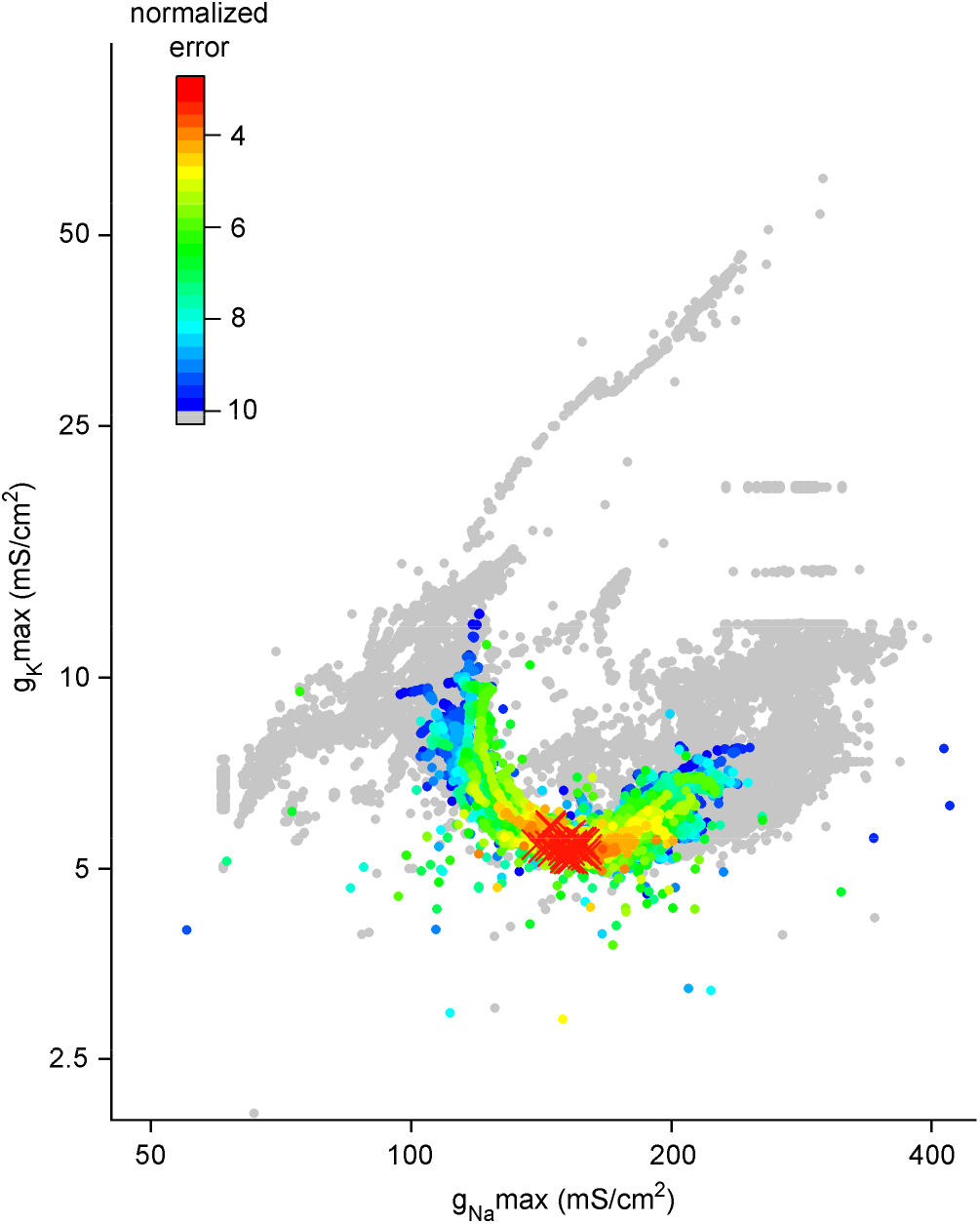
Supporting data for Figure 4. Uniqueness of the fit results. The explored parameter space during the optimization of the conductances needed to reconstruct the recorded APs (30 fitted APs started from 120 initializations). Each dot represent single run during the fit (n=142605 runs). The optimization error of the each run is color coded. Red crosses mark best-fit solutions.

**Supplementary Figure 4.**
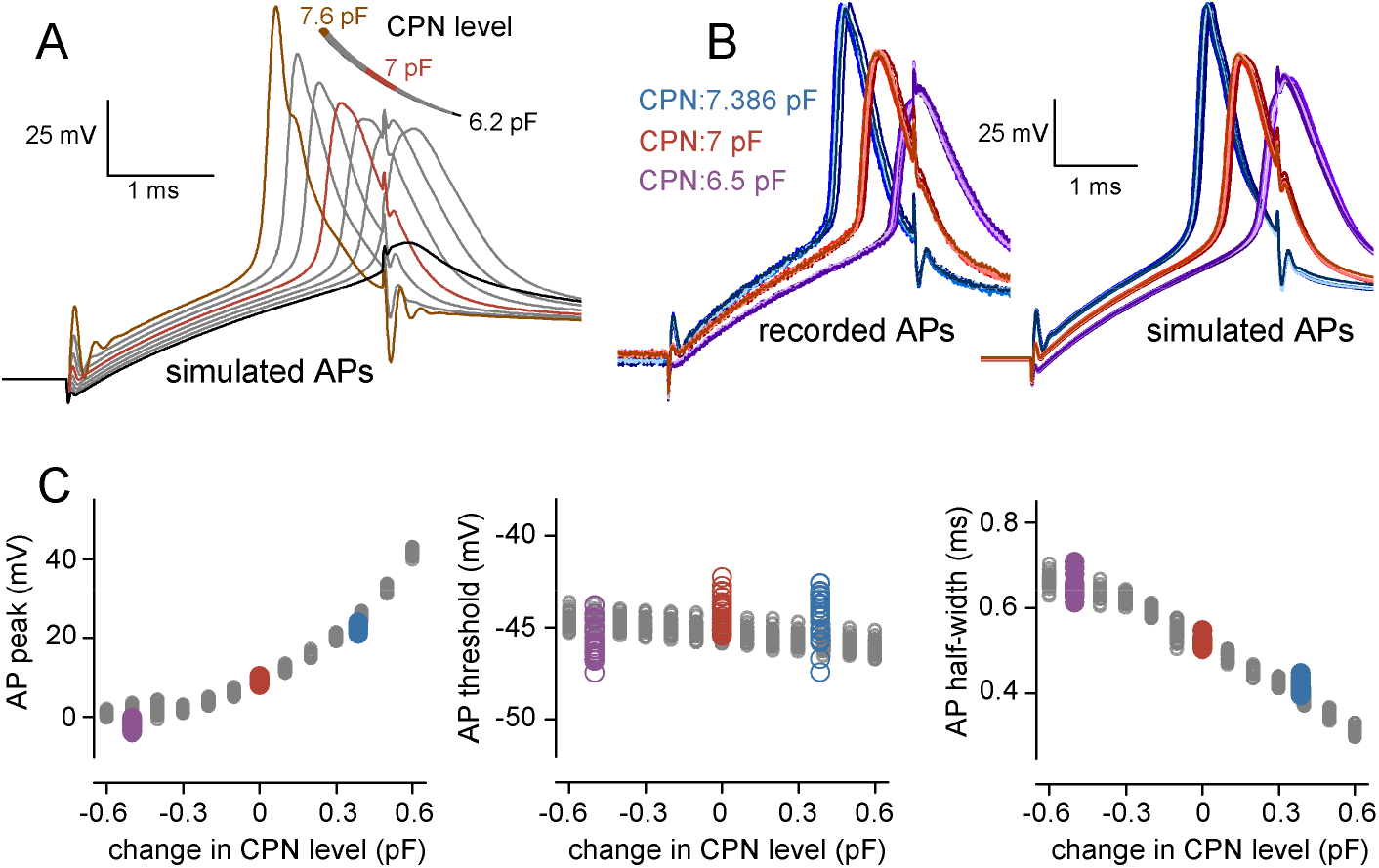
Supporting data for Figure 5. Critical influence of the applied CPN on the recorded AP waveforms. (A) Model AP waveforms simulated using the same conductance set but different CPN settings. Notice the AP failure at CPN=6.2 pF (black) and the oscillation at CPN=7.6 pF (brown). (B) Representative APs recorded (left) and simulated (right) in the same axon with different CPN settings (n=6 APs in each CPN conditions). (C) Effects of different CPN level on AP peak (left), threshold (middle) and half-width (right). Gray circles show the AP parameters simulated with the 30 different conductance sets (the same set as in **Figure 4**). Purple, red and blue symbols show the experimentally recorded AP parameters obtained with three different CPN settings (n=30 APs in each conditions). Zero on x-axis represents the originally set 7 pF capacitance neutralization.

**Supplementary Figure 5.**
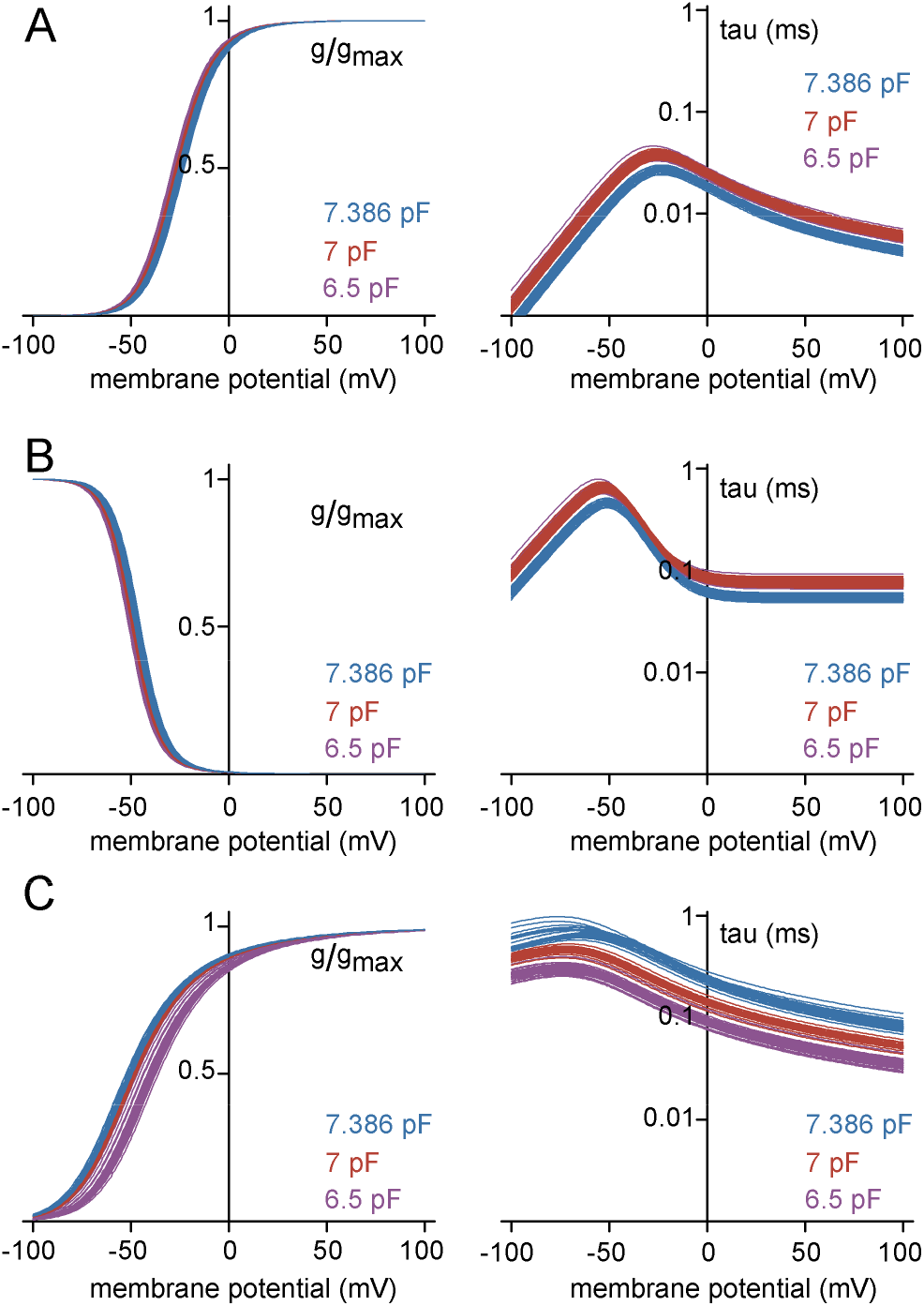
Supporting data for Figure 5. Gating profile of model conductances obtained from fitting of AP waveforms with different levels of distortions. (A) Voltage dependence (left) and kinetic profile (right) of the activation in Na^+^ conductance models obtained by the reconstruction of the APs recorded with different CPN settings (CPN = 6.5 pF in purple, CPN = 7 pF in red and CPN = 7.386 pF in blue). Each line represent single model conductance (n=30/30/28 models). (B) Same as in panel (A) but for the inactivation of the Na^+^ conductance models. (C) Same as in panel (A) but for the K^+^ conductance models

**Supplementary Figure 6.**
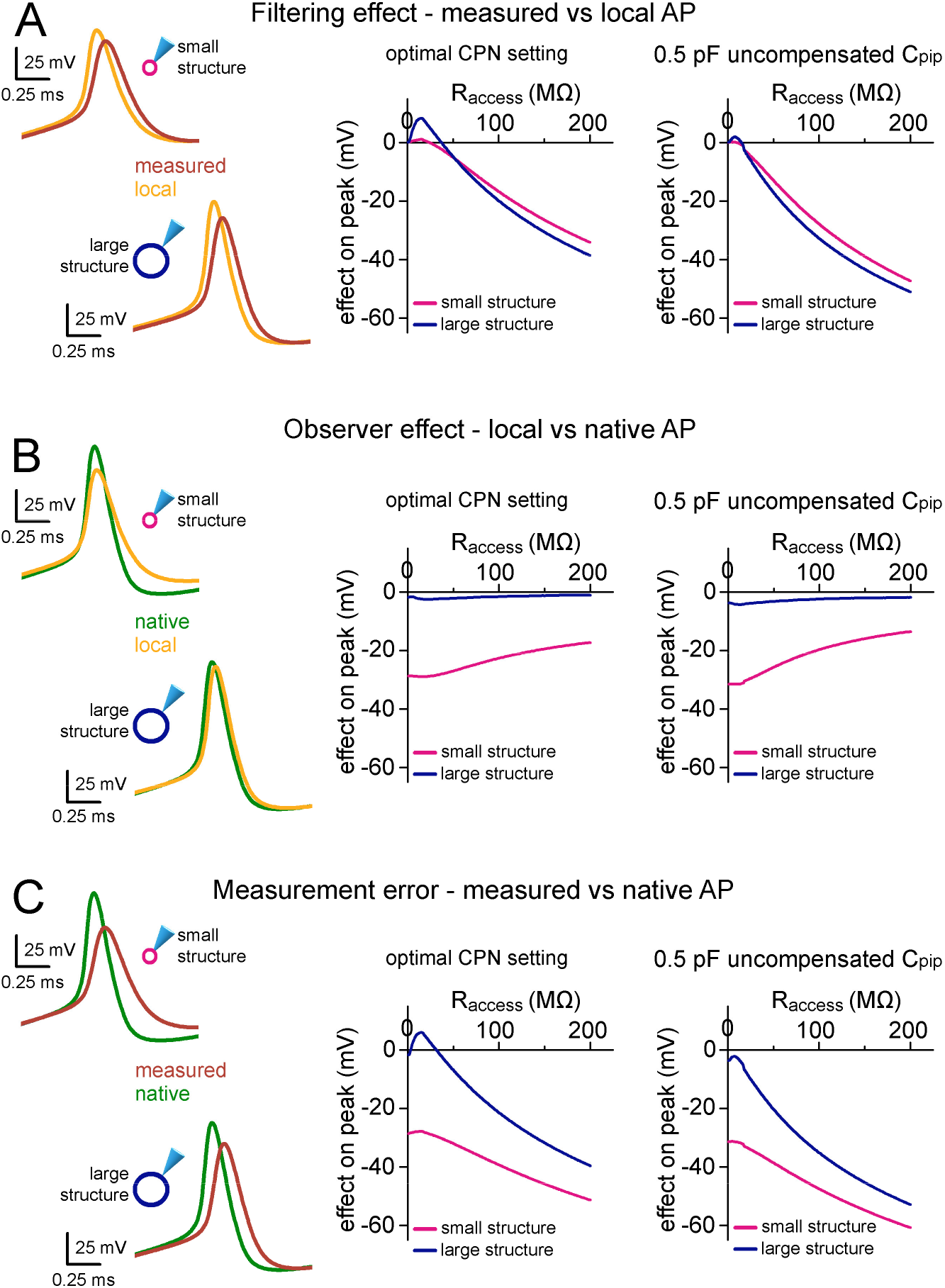
Supporting data for Figure 7. Instrumental and structural parameters cooperatively determine signal distortions in recordings from small neuronal structures. The results of the same simulations as in **Figure 7**, are shown for effects on AP peak.

